# Investigating Motor Skill Acquisition in Mice and Humans: Towards A Cross-Species Paradigm

**DOI:** 10.64898/2026.11.20.768070

**Authors:** Maike Hille, Lena C. Justus, Sara Zocher, Tibor Stark, Moritz Bonhoeffer, Jason P. Lerch, Michael Czisch, Simone Kühn, Ulman Lindenberger, Tobias Bonhoeffer

**Author notes:** shared first authorship (Maike Hille, Lena C. Justus, Sara Zocher, Tibor Stark). shared senior authorship (Simone Kühn, Ulman Lindenberger, Tobias Bonhoeffer).

## Abstract

Relating non-invasive observations of human brain plasticity to cellular processes identified in animal models remains a major challenge in neuroscience. We address this challenge by introducing a paradigm for studying motor skill acquisition across species. We designed a human chopstick grasping task modeled after the rodent single-pellet reaching task, resulting in a comparable sequence of reach-to-grasp movements across species. We studied 48 young adult male C57BL/6N mice and 71 young men, comparing trained groups with untrained controls. In both mice and humans, structural magnetic resonance imaging (MRI) was performed before, during, and after motor training. The functional form of behavioral performance gains on the trained task was highly similar across species. In trained compared to control mice, analysis of the early learning interval revealed decreases in regional volume in contralateral primary motor and forelimb somatosensory cortices, with greater volume reductions in primary motor cortex associated with better reaching performance. In humans, performance gains transferred to other chopstick tasks but not to general dexterity, and no reliable changes in gray matter volume were observed. Quantitative MRI, however, revealed early training-related changes in tissue-sensitive measures, including increases in magnetization transfer saturation (MT_sat_) and longitudinal relaxation rate (R_1_) in contralateral motor and frontal regions, potentially reflecting an increase in intracortical myelin. Thus, early structural changes in the contralateral motor cortex were detected in both species, but the degree to which these changes reflect a common biological mechanism is unclear. We conclude that the present cross-species paradigm provides a viable starting point for studying brain plasticity across species. Future work needs to relate macroscopic structural changes more directly to cellular processes by applying longitudinal microscopic imaging and MRI in the same animals.

## Introduction

Understanding the mechanisms underlying human brain function is a central goal of neuroscience. It ultimately requires the characterization of the processes occurring in the brain’s cells and synapses, a level of detail that remains largely inaccessible in living humans. Much of this knowledge therefore comes from animal models, which have provided fundamental insights into these cellular and molecular processes. Applying these insights to humans rests on the premise that mechanisms identified in animals generalize across species, an assumption that is rarely tested directly. Addressing this gap requires study designs that can be implemented in comparable form across species, where outcomes can be measured with the same method – a combination few experimental paradigms offer (Hille et al., 2024). Motor skill acquisition meets both requirements: it is a well-established experimental model of experience-dependent plasticity in humans and rodents, and its structural consequences can be measured with magnetic resonance imaging (MRI) in both species.

In humans, longitudinal structural MRI (sMRI) studies have shown that motor skill acquisition is associated with regional changes in morphometric measures, such as gray matter volume. Training-related gray matter changes were first reported after juggling (Draganski et al., 2004; Driemeyer et al., 2008; Scholz et al., 2009), and subsequently for whole-body balance training, golf training, and non-dominant hand motor practice, implicating distributed sensorimotor, premotor, parietal, and cerebellar regions in experience-dependent structural plasticity during motor learning (Bezzola et al., 2011; Gryga et al., 2012; Taubert et al., 2010; Wenger, Kühn, et al., 2017). Importantly, these changes may not necessarily follow a monotonic trajectory: volume may increase and renormalize during continued training, consistent with an expansion–renormalization account of brain plasticity (Lindenberger & Lövdén, 2019; Lövdén et al., 2013, 2020; Wenger, Brozzoli, et al., 2017). Furthermore, changes can emerge rapidly, even within a single training session (Kodama et al., 2018; Olivo et al., 2022; Taubert et al., 2016). However, these macroscopic changes are difficult to interpret, as they are biologically unspecific and cannot, on their own, identify the underlying cellular mechanisms (Hille et al., 2024; Zatorre et al., 2012). Quantitative MRI (qMRI) has therefore been introduced alongside conventional sMRI to obtain measures more closely related to tissue microstructure and composition (Weiskopf et al., 2021). During motor learning, changes in several qMRI measures, including magnetization transfer saturation (MT_sat_) and longitudinal relaxation rate (R_1_) have been reported, with MT_sat_ and R_1_ being interpreted to be associated with changes in myelin content and remodeling given their sensitivity to macromolecular content and tissue composition (Aye et al., 2025; Azzarito et al., 2023).

Still, these measures remain indirect indicators of the cellular mechanisms underlying experience-dependent plasticity. By contrast, histological studies and longitudinal multiphoton microscopy in rodents have revealed several cellular processes associated with motor skill acquisition, including changes in synaptic connectivity (Fu et al., 2012; Fu & Zuo, 2011; Xu et al., 2009; Yang et al., 2009) and intracortical myelination (Bacmeister et al., 2020; McKenzie et al., 2014; Sampaio-Baptista et al., 2013). Applying sMRI in rodents could therefore offer an important intermediate level of analysis between cellular mechanisms and human neuroimaging findings (Hille et al., 2024). However, only few studies have made that link, and those that have done so reported both gray matter increases and decreases across different paradigms and brain regions (Badea et al., 2019; Biedermann et al., 2016; Lerch et al., 2011; Mediavilla et al., 2022; Scholz, Allemang-Grand, et al., 2015; Scholz, Niibori, et al., 2015).

Among these, the single-pellet reaching task, which we selected for our cross-species comparison, illustrates the general difficulty of interpreting macroscopic changes. In one study, gray matter volume in the motor cortex increased after 25 days of training (Badea et al., 2019), whereas another study found a decrease over 15 days of training (Mediavilla et al., 2022). Notably, in the latter, the decrease was accompanied by increases in white matter volume within the same cortical regions, and the authors attributed both changes to increased intracortical myelin rather than tissue loss. Even when sMRI and multiphoton microscopy were combined in the same animals, albeit focusing on aging not intervention effects, only a third of the variance in gray matter volume could be explained by physical volume change, cell density and nucleus volume together (Asan et al., 2021).

Despite these advances, the requirements for a direct comparison across species are currently not met: Human and rodent studies typically use tasks that differ substantially in motor demands, training duration, and behavioral outcome measures. Consequently, it cannot be determined whether observed neural changes reflect shared principles of skill learning or species- and task-specific effects. A suitable paradigm that combines longitudinal sMRI with microscopic measures in rodents would provide a methodological interface to relate rodent cellular measures to human MRI findings and thereby improve their biological interpretability (Hille et al., 2024).

Skilled reaching is a natural candidate for such a paradigm: It involves a comparable sequence of reach-to-grasp movements in humans and rodents (Sacrey et al., 2009), and engages sensorimotor cortical regions that are likely homologous across species (Bakken et al., 2021; BRAIN Initiative Cell Census Network (BICCN) et al., 2021; Lazari et al., 2024). We therefore developed a human grasping task adapted from the single-pellet reaching task in rodents, in which participants used chopsticks to grasp individual roundish sweets through a narrow slit (Figure 1). Cohorts of mice and humans were trained on the respective tasks to assess whether they showed comparable trajectories of motor performance improvements. Both species were scanned longitudinally using sMRI to obtain morphometric measures and assess whether training induced comparable macroscopic structural changes in humans and mice. We expected both species to improve in performance with training and to show similar learning trajectories, with the largest gains occurring early in training and decreasing over time. Based on previous longitudinal studies of fine-motor skill learning, we hypothesized that training would induce a curvilinear trajectory of gray matter volume in contralateral primary motor cortex, characterized by an initial increase followed by a decrease over the course of training (Wenger, Kühn, et al., 2017). We expected a comparable pattern of change in regional volume in mice, reflecting similar macroscopic structural dynamics during motor skill training. In humans, we additionally obtained qMRI parameters and analyzed them in an exploratory manner to provide further insight into tissue properties that might underlie any observed volumetric changes.

**Figure 1.**
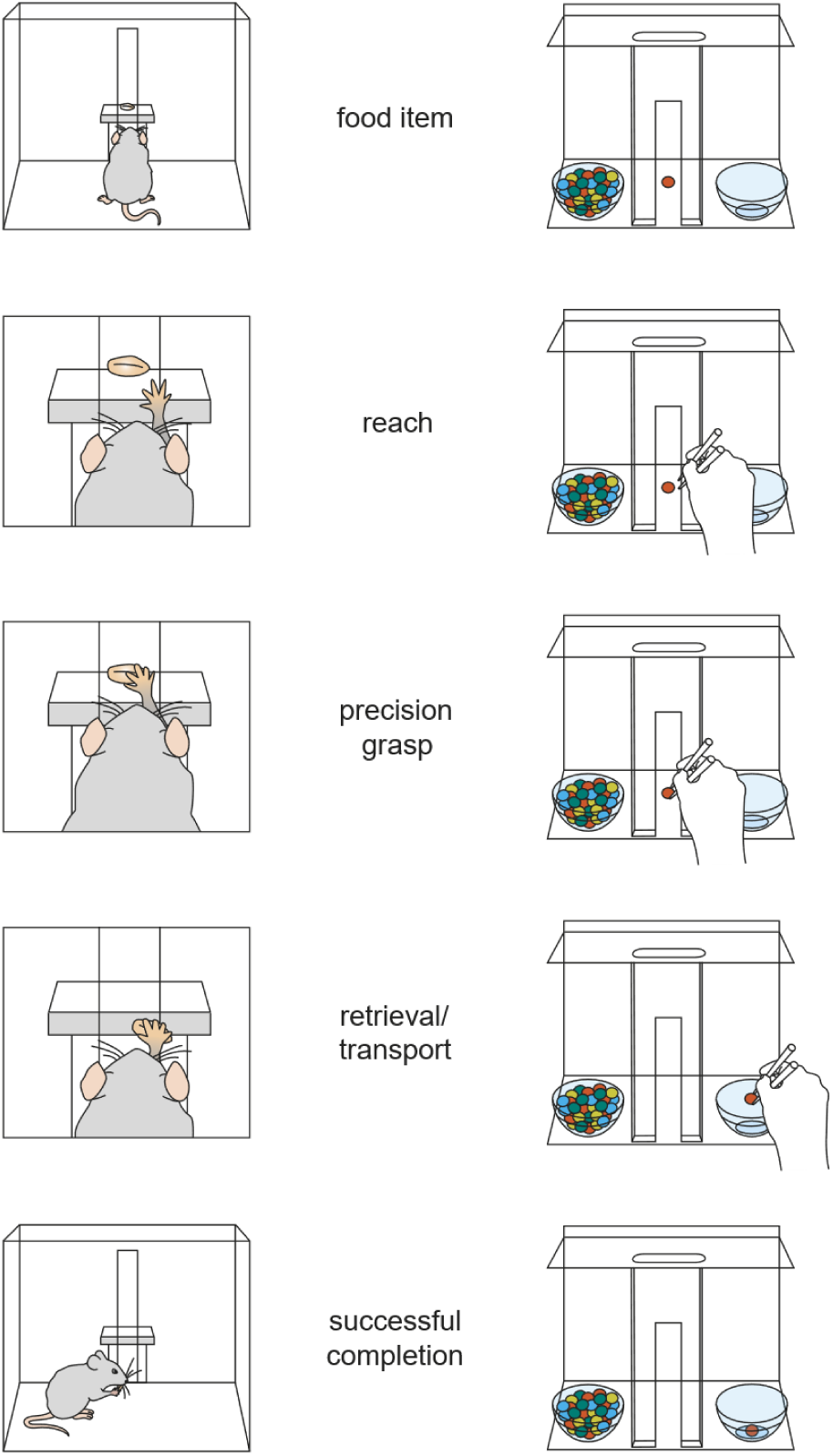
Schematic illustration of the mouse (left) and the human (right) grasping tasks. Both tasks follow a comparable sequence comprising presentation of a food item, reaching, precision grasping, retrieval and transport of the item, and successful task completion.

## Results

### Motor skill acquisition paradigm

In the rodent single-pellet reaching task, animals perform precise forelimb reach-to-grasp movements to retrieve individual millet seeds through a narrow slit. We adapted this structure for humans: participants used a pair of chopsticks to grasp single M&Ms (chocolate-coated peanuts) through a slit and transfer them to a bowl (Figure 1). In both species, each session comprised 30 trials, analyzed in three bins of ten. Mice consumed each millet seed immediately after grasping, whereas human participants were allowed to consume the sweets after task completion. Performance was quantified as success rate in mice and as grasping time in humans. Mice were assigned either to a forelimb training group, or a control group that received identical handling and were put inside the grasping box without undergoing specific training. Human participants were assigned to a chopstick training or a reading control group. Mice were trained daily except for MRI acquisition days, for eight sessions over 11 days (Figure 2A and Figure S2). Human participants practiced daily over 70 days, supervised on MRI acquisition days and otherwise at home (Figure 2A). Individual performance trajectories across the at-home training period are shown in Figure S2. The seven supervised sessions (S1–S7) on-site provided the measurements reported below.

**Figure 2.**
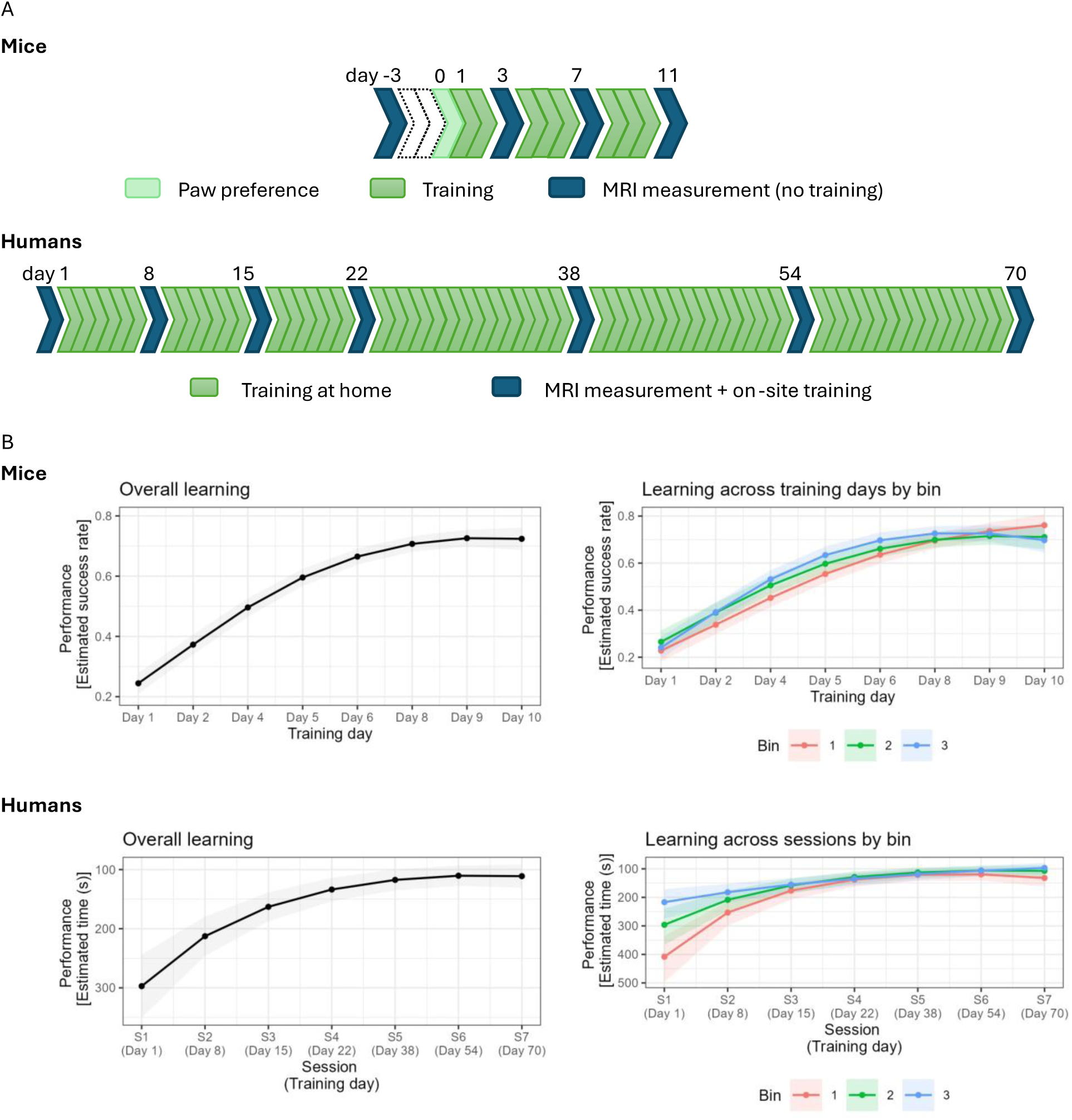
***A,*** Experimental design for mice (top) and humans (bottom). The mouse study comprised an 11-day training period, preceded by a baseline MRI measurement on day −3 and paw preference testing on day 0. Further MRI measurements were acquired on days 3, 7, and 11. Mice underwent single-pellet reaching training on the intervening non-scan days, beginning on day 1. The human study followed a 70-day longitudinal design with seven in-person sessions on days 1 (baseline), 8, 15, 22, 38, 54, and 70. MRI measurements were acquired at each session. Between sessions, participants performed daily chopstick grasping training or a matched-duration reading control task. ***B,*** Behavioral learning curves for mice (top, *n* = 32) and humans (bottom, *n* = 36). Left panels show overall learning across training, based on model-estimated performance averaged across the three within-session bins. Right panels show model-estimated performance separately for each bin. Points and lines indicate model-estimated marginal means, and shaded areas indicate 95% confidence intervals. Mouse performance was quantified as reach success probability, with higher values indicating better performance; bins 1–3 correspond to reaches 1–10, 11–20, and 21–30, respectively. Human performance was quantified as grasping time, with lower values indicating better performance; the y-axis is plotted in reverse order so that improved performance is displayed upward. Bins 1–3 correspond to M&Ms 1–10, 11–20, and 21–30, respectively.

#### Mice showed progressive improvement in skilled reaching across training

For the mice (*n* = 32), we examined learning curves by fitting a generalized linear mixed-effects model predicting the number of successful reaches out of 10 attempts per bin as a function of training day (linear and quadratic terms, mean-centered), bin (reaches 1–10, 11–20, 21–30), and their interactions, with mouse-specific random intercepts and random slopes for training day per mouse (Figure 2B). The model revealed significant main effects of both the linear (χ²(1) = 185.14, *p* < .001) and quadratic (χ²(1) = 12.89, *p* < .001) training day terms, indicating a curvilinear improvement in reach success across training days. There was also a significant main effect of bin (χ²(2) = 12.13, *p* = .002), as well as significant linear (χ²(2) = 6.71, *p* = .035) and quadratic (χ²(2) = 7.07, *p* = .029) Training Day × Bin interactions, indicating that both the overall level of performance and the shape of the learning curve differed across bins.

We compared model-estimated reach success probabilities between consecutive training days, averaged across bins to better characterize the overall pattern of improvement across training (Table 1). Success probability increased steadily between consecutive training days from training day 1 to training day 9 with the magnitude of improvement progressively decreasing across training. No significant change was observed between training days 9 and 10, suggesting that mice approached asymptotic performance by training day 9. These results show that improvement was most pronounced early in training and progressively decelerated, plateauing by the final training days.

**Table 1.** Pairwise comparisons of mice grasping performance across training days.

| **Training day comparison** | **Estimated grasping success (%)** | **Change (pp)** | ***z*** | ***p*** |
|---|---|---|---|---|
| Day 1 → Day 2 | 24.4 → 37.3 | 12.8 | 19.8 | < .001 |
| Day 2 → Day 4 | 37.3 → 49.6 | 12.3 | 18.1 | < .001 |
| Day 4 → Day 5 | 49.6 → 59.6 | 9.9 | 18.1 | < .001 |
| Day 5 → Day 6 | 59.6 → 66.5 | 7.0 | 15.4 | < .001 |
| Day 6 → Day 8 | 66.5 → 70.7 | 4.2 | 9.0 | < .001 |
| Day 8 → Day 9 | 70.7 → 72.6 | 1.9 | 3.3 | .002 |
| Day 9 → Day 10 | 72.6 → 72.4 | −0.2 | −0.3 | .787 |

*Note.* Estimated success probabilities are model-estimated marginal means averaged across bins. Change is expressed in percentage points (pp) relative to the preceding training day, with positive values indicating increased reach success. *p* values were Holm-adjusted across all seven comparisons between consecutive assessed training days.

We followed up on the significant Training Day × Bin interaction by comparing reach success probability between consecutive bins (bin 2 vs. bin 1; bin 3 vs. bin 2) separately for each training day (Table 2). Although the pattern of performance across bins varied over the course of training, none of the individual consecutive-bin comparisons remained significant after Holm correction (all *p*s ≥ .116).

**Table 2.** Pairwise comparisons of mouse grasping performance between consecutive bins within training days.

| **Training day** | **Bin comparison** | **Estimated grasping success (%)** | **Change (pp)** | ***z*** | ***p*** |
|---|---|---|---|---|---|
| Day 1 | Bin 1 → Bin 2 | 22.8 → 26.5 | 3.7 | 1.34 | 1.000 |
|  | Bin 2 → Bin 3 | 26.5 → 24.1 | −2.4 | −0.86 | 1.000 |
| Day 2 | Bin 1 → Bin 2 | 33.8 → 39.0 | 5.1 | 2.42 | .232 |
|  | Bin 2 → Bin 3 | 39.0 → 39.2 | 0.2 | 0.10 | 1.000 |
| Day 4 | Bin 1 → Bin 2 | 45.2 → 50.5 | 5.3 | 2.68 | .116 |
|  | Bin 2 → Bin 3 | 50.5 → 53.1 | 2.6 | 1.30 | 1.000 |
| Day 5 | Bin 1 → Bin 2 | 55.4 → 59.7 | 4.3 | 2.03 | .594 |
|  | Bin 2 → Bin 3 | 59.7 → 63.4 | 3.7 | 1.77 | .930 |
| Day 6 | Bin 1 → Bin 2 | 63.6 → 66.1 | 2.5 | 1.26 | 1.000 |
|  | Bin 2 → Bin 3 | 66.1 → 69.7 | 3.6 | 1.85 | .841 |
| Day 8 | Bin 1 → Bin 2 | 69.6 → 69.9 | 0.4 | 0.21 | 1.000 |
|  | Bin 2 → Bin 3 | 69.9 → 72.6 | 2.7 | 1.59 | 1.000 |
| Day 9 | Bin 1 → Bin 2 | 73.6 → 71.5 | −2.1 | −1.13 | 1.000 |
|  | Bin 2 → Bin 3 | 71.5 → 72.6 | 1.1 | 0.60 | 1.000 |
| Day 10 | Bin 1 → Bin 2 | 76.1 → 71.1 | −5.0 | −1.72 | .939 |
|  | Bin 2 → Bin 3 | 71.1 → 69.8 | −1.3 | −0.43 | 1.000 |

*Note.* Estimated grasping success probabilities are back-transformed estimated marginal means from the generalized linear mixed-effects model. Bin 1, Bin 2, and Bin 3 correspond to reaches 1–10, 11–20, and 21–30, respectively. Changes are expressed in percentage points (pp), calculated from unrounded estimates relative to the preceding bin, with positive values indicating increased grasping success. *p* values were Holm-adjusted across all 16 comparisons between consecutive bins.

#### Human chopstick grasping training produced a task-specific motor skill

To assess performance in humans following training with the newly developed chopstick grasping task, we first compared the chopstick training group (*n* = 33) with the reading control group (*n* = 32) at the first (S1) and final (S7) sessions. Performance on the trained chopstick grasping task improved more in the chopstick training group than in the reading control group from S1 to S7 (Figure S1, Table S1). Mixed-effects models showed a significant Session × Group interaction for the 30 M&Ms task (*F*(1,63) = 67.27, *p* < .001). Completion times decreased significantly from 1040 s at S1 to 315 s at S7 in the chopstick training group, corresponding to a 69.7% reduction (*t*(63) = −15.30, *p* < .001). Completion times in the reading control group decreased significantly less, from 1008 s to 760 s (24.6% reduction, *t*(63) = −3.56, *p* < .001). At S1, the groups did not differ in completion time (*t*(115) = 0.34, *p* = .735), whereas at S7, the chopstick training group completed the task 58.5% faster than the reading control group did (*t*(115) = −9.34, *p* < .001).

Similar training-related improvements were also observed across other chopstick grasping tasks (near transfer), including tasks involving different objects and use of the non-dominant hand (Table S1). In contrast, we found no evidence that chopstick training altered performance on the more general fine motor dexterity measures (far transfer). Neither the Purdue Pegboard, the Grooved Pegboard, nor the O’Connor Tweezer Dexterity test showed a significant Session × Group interaction (Table S2). Importantly, we additionally confirmed that groups did not differ significantly at S1 on any of these measures, indicating a comparable baseline of fine motor dexterity across the groups.

To characterize changes in performance over the course of training, we analyzed session-wise learning curves in the chopstick training group (*n* = 36). For the human participants, we fitted a linear mixed-effects model predicting log-transformed grasping time, with fixed effects of session (linear and quadratic terms, mean-centered), bin (M&Ms 1–10, 11–20, 21–30), and their interactions, as well as random intercepts for participants, and random slopes for session per participant (Figure 2B). The model revealed significant main effects of both the linear (*F*(1, 34.1) = 106.35, *p* < .001) and quadratic (*F*(1, 591.4) = 47.24, *p* < .001) session terms. There was no significant main effect of bin (*F*(2, 577.5) = 0.69, *p* = .504). The linear session × bin interaction was not significant (*F*(2, 582.9) = 2.28, *p* = .103), whereas the quadratic session × bin interaction was significant (*F*(2, 581.3) = 8.41, *p* < .001), indicating that the shape of the learning curve differed across bins.

To further characterize the overall pattern of improvement across sessions, we compared model-estimated grasping times between consecutive sessions, averaged across bins (Table 3). Grasping time decreased significantly between all consecutive sessions from S1 to S6, with the magnitude of improvement progressively decreasing across training. No significant change was observed between S6 and S7, suggesting that participants were close to asymptotic performance by S6. These results show that improvement was most pronounced early in training and progressively decelerated, plateauing by the final sessions, a pattern paralleling the learning trajectory observed in mice.

**Table 3.** Pairwise comparisons of human grasping performance across training sessions.

| **Session comparison** | **Estimated grasping time (s)** | **Change (%)** | ***t*** | ***df*** | ***p*** |
|---|---|---|---|---|---|
| S1 → S2 | 297.2 → 212.7 | −28.4 | −11.07 | 340.2 | < .001 |
| S2 → S3 | 212.7 → 163.0 | −23.4 | −11.98 | 141.8 | < .001 |
| S3 → S4 | 163.0 → 133.7 | −18.0 | −12.01 | 46.5 | < .001 |
| S4 → S5 | 133.7 → 117.4 | −12.2 | −8.26 | 37.8 | < .001 |
| S5 → S6 | 117.4 → 110.4 | −6.0 | −3.02 | 101.1 | .038 |
| S6 → S7 | 110.4 → 111.1 | 0.6 | 0.23 | 269.9 | 1.000 |

*Note.* Estimated grasping times are back-transformed estimated marginal means from the linear mixed-effects model and are averaged across bins. Change (%) is calculated relative to the earlier session, with negative values indicating a reduction in grasping time. *p* values are Holm-adjusted for multiple comparisons.

To decompose the significant quadratic Session × Bin interaction, we compared performance between consecutive bins (bin 2 vs. bin 1; bin 3 vs. bin 2) separately at each session (Table 4). Within-session improvements were most pronounced early in training, with significant decreases in grasping time from bin 1 to bin 2 at S1 and S2. No other consecutive-bin comparisons were significant, consistent with a within-session practice effect that was strongest before participants approached asymptotic performance.

**Table 4.** Pairwise comparisons of human grasping performance between consecutive bins within training sessions.

| **Session** | **Bin comparison** | **Estimated grasping time (s)** | **Change (%)** | ***t*** | ***df*** | ***p*** |
|---|---|---|---|---|---|---|
| S1 | Bin 1 → Bin 2 | 408.3 → 296.2 | −27.5 | −3.37 | 589.1 | .010 |
|  | Bin 2 → Bin 3 | 296.2 → 217.0 | −26.8 | −2.76 | 583.5 | .071 |
| S2 | Bin 1 → Bin 2 | 253.3 → 208.7 | −17.6 | −3.42 | 586.4 | .009 |
|  | Bin 2 → Bin 3 | 208.7 → 182.0 | −12.8 | −2.14 | 582.0 | .305 |
| S3 | Bin 1 → Bin 2 | 176.4 → 158.1 | −10.4 | −2.01 | 578.2 | .359 |
|  | Bin 2 → Bin 3 | 158.1 → 155.2 | −1.8 | −0.32 | 577.5 | 1.000 |
| S4 | Bin 1 → Bin 2 | 137.9 → 128.7 | −6.7 | −1.17 | 577.2 | 1.000 |
|  | Bin 2 → Bin 3 | 128.7 → 134.6 | 4.6 | 0.73 | 577.2 | 1.000 |
| S5 | Bin 1 → Bin 2 | 121.1 → 112.6 | −7.0 | −1.31 | 577.6 | 1.000 |
|  | Bin 2 → Bin 3 | 112.6 → 118.7 | 5.4 | 0.91 | 577.6 | 1.000 |
| S6 | Bin 1 → Bin 2 | 119.4 → 105.9 | −11.2 | −2.17 | 576.7 | .305 |
|  | Bin 2 → Bin 3 | 105.9 → 106.4 | 0.4 | 0.07 | 576.8 | 1.000 |
| S7 | Bin 1 → Bin 2 | 132.1 → 107.1 | −18.9 | −2.35 | 576.6 | .211 |
|  | Bin 2 → Bin 3 | 107.1 → 96.9 | −9.5 | −1.11 | 576.6 | 1.000 |

***Note.*** Bin 1, Bin 2, and Bin 3 correspond to M&Ms 1–10, 11–20, and 21–30, respectively. Statistical tests were performed on the log scale; estimated grasping times are expressed in seconds. Percentage changes were calculated from unrounded estimates relative to the preceding bin, with negative values indicating a reduction in grasping time. *p* values were Holm-adjusted across all 14 comparisons between consecutive bins.

#### Learning trajectories were comparable across species

To quantitatively compare the shapes of the learning trajectories across species, we fitted separate exponential learning functions to the population-level performance data for humans and mice. In both species, performance followed the characteristic pattern of rapid improvement early in training followed by progressively smaller gains as performance approached an asymptote. The estimated learning rate parameter was k = 0.31 (95% CI [0.23, 0.38]) in humans and k = 0.32 (95% CI [0.24, 0.40]) in mice, indicating a similar rate of convergence toward asymptotic performance across sessions in the two species.

### Training-related structural brain changes

To assess training-related structural brain changes in both species we acquired MRI at multiple timepoints across learning. In mice structural images were acquired for atlas-based segmentation of regional volumes while in human participants we collected structural images for voxel-based morphometry (VBM) and qMRI parameter maps. The timing of scans was designed to cover both initial rapid plasticity and slower expansion–renormalization patterns.

#### Sensorimotor cortical volume decreased relative to controls during early learning in mice

Before examining training-related changes in regional volume in mice, we first assessed total brain volume of control (*n* = 16) and trained mice (*n* = 17; 16 on day 11, as one scan failed registration) over the imaging period. At baseline, we found comparable volumes in control and trained mice (control *M* = 409.6 mm^3^, *SD* = 10.3, trained *M* = 406.1 mm^3^, *SD* = 9.5; Student’s *t*(31) = −1.03, *p* = .31). The total brain volume decreased in both groups over the four MRI scans (−1.3% in control and −1.0% in trained animals; linear mixed model, main effect of time *F*(3,92.0) = 32.87, *p* < .001; Figure S3). This decrease did not differ between control and trained mice (Time × Group interaction *F*(3,92.0) = 0.75, *p* = .53). The anesthesia control group, trained identically but only scanned twice (S1 [day −3] and S4 [day 11], *n* = 13), showed a smaller decrease (−0.7%), which did not differ significantly from that of trained mice scanned four times (Welch’s *t*(18.5) = 0.86, *p* = .40; Figure S3). Total brain volume was included as a covariate in all region-of-interest (ROI) group comparisons.

We restricted the analysis to primary motor cortex and forelimb somatosensory cortex, both contralateral to the trained paw. Using a linear mixed model across all four timepoints we detected no significant Time × Group interaction for either the primary motor (*F*(3,84.0) = 1.51, *p* = .22, *p*_Holm_ = .22) or forelimb somatosensory cortex (*F*(3,90.9) = 2.57, *p* = .059, *p*_Holm_ = .12). The primary comparison of the early learning period between S1 and S2 (after two days of training) showed that the change in regional volume was more negative in trained than in control animals in both regions (Figure 3A): primary motor cortex (estimate −0.071 mm^3^, 95% CI [−0.140, −0.003]; *t*(84.2) = −2.06, *p* = .042, *p*_Holm_ = .042) and forelimb somatosensory cortex (estimate −0.046 mm^3^, 95% CI [−0.080, −0.012]; *t*(91.1) = −2.71, *p* = .008, *p*_Holm_ = .016). Follow-up within-group contrasts from the respective models showed that volume decreased significantly in trained mice in forelimb somatosensory cortex (−2.5%, *p* < .001, *p*_Holm_ < .001, Table S3), but not in primary motor cortex (−1.3%, *p* = .067, *p*_Holm_ = .20). No significant volume changes were observed in control animals in either region (primary motor +0.7 %, *p* = .33, *p*_Holm_ = .66; forelimb somatosensory −0.1%, *p* = .86, *p*_Holm_ = .86; Table S3).

**Figure 3.**
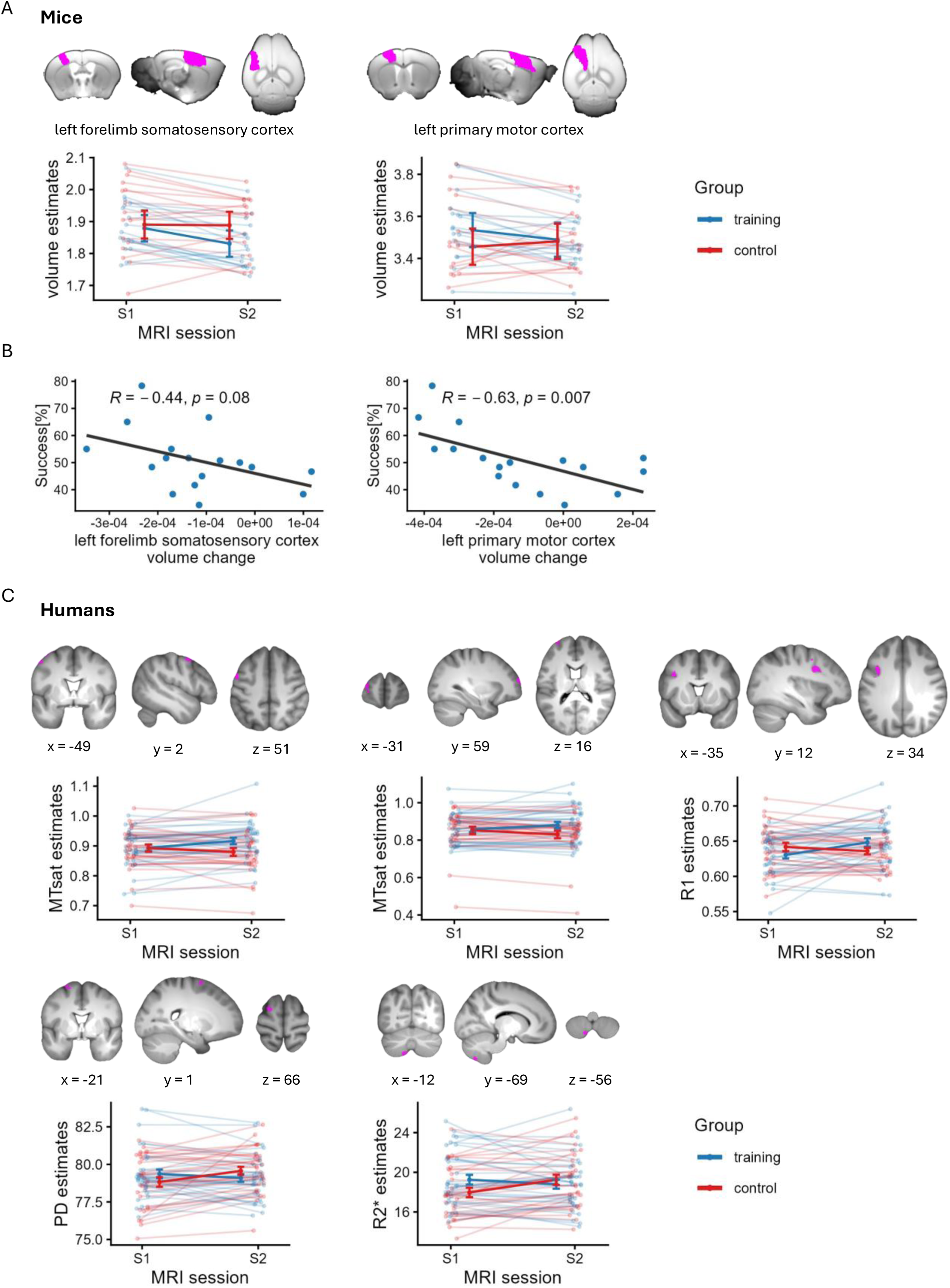
***A,*** Changes in regional volume from S1 (day −3) to S2 (day 3) in mice. Thin lines represent individual mouse trajectories, while thick lines connect estimated marginal means derived from linear mixed-effects models adjusted for total brain volume, with a random intercept for each mouse. Error bars indicate 95% confidence intervals. Training group *n* = 17, control group *n* = 16. ***B,*** Correlation between observed volume change from S1 to S2 and behavioral performance averaged across training days 2 (before S2) and 4 (after S2) in mice. ***C,*** Changes in qMRI parameter maps from session 1 to session 2 in humans (*p* < .001 uncorrected, cluster-size threshold k = 232 voxels). Thin lines represent individual human participant trajectories, while thick lines connect observed group means across sessions. Data were extracted from significant clusters identified in the whole-brain analysis and are presented for descriptive purposes only. Error bars indicate ±1 standard error of the mean (SEM). Training group *n* = 33, control group *n* = 30.

#### Motor cortical volume change correlated with reaching performance in mice

To assess if volumetric changes of the analyzed brain regions were associated with task performance, we further correlated the change in regional volume relative to total brain volume early in training with behavioral performance in trained mice (*n* = 17), measured as the mean success rate across the two full training sessions flanking the second scan (Figure 3B). A greater volume decrease in primary motor cortex correlated with better reaching performance (Pearson’s *r* = −0.63, *p* = .007, *p*_Holm_ = .014, 95% CI [−0.85, −0.21]), whereas the correlation in forelimb somatosensory cortex showed a similar negative association but did not reach significance (*r* = −0.44, *p* = .080, *p*_Holm_ = .080, 95% CI [−0.76, +0.06]). Rank correlations were larger in magnitude in both regions (primary motor cortex Spearman’s *r*_s_ = −0.70, *p* = .004, *p*_Holm_ = .008, 95% CI [−0.89, −0.27]; forelimb somatosensory cortex *r*_s_ = −0.49, *p* = .056, *p*_Holm_ = .056, 95% CI [−0.80, +0.01]). The raw change in total brain volume over the same interval did not correlate with behavioral performance (Pearson’s *r* = −0.23, *p* = .38, 95% CI [−0.64, +0.29], uncorrected).

#### VBM-based analyses did not show training-related gray matter volume changes in humans

For the human participants, across all MRI analyses of the VBM-processed structural T1-weighted images, we did not detect any significant training-related gray matter volume changes. Specifically, the whole-brain flexible factorial model did not reveal any significant Time × Group interactions. Furthermore, the ROI-based analyses of gray matter volume estimates extracted from task-defined motor regions showed no significant Time × Group interactions. The exploratory analysis based on individual learning curves did not show a significant effect within the training group either.

#### Quantitative MRI parameters showed training-related changes across human cortical and cerebellar regions

Whole-brain analyses of the four qMRI parameter maps, MT_sat_, R_1_, PD (proton density) and effective transverse relaxation rate (R_2_*), revealed significant Time × Group interactions in several clusters, surviving the predefined voxel-wise threshold of *p* < .001 uncorrected and the cluster-extent thresholds of k ≥ 232 voxels (Figure 3C and Figure S4). Specifically, for the contrast testing a greater S1-to-S2 increase in the training group than in the control group, significant clusters were observed for MT_sat_ in the premotor cortex, encompassing the left precentral and left middle frontal gyrus (x = −49, y = 2, z = 51; k = 407 voxels) and in the left middle frontal gyrus (x = −31, y = 59, z = 16; k = 328 voxels). Furthermore, we observed significant clusters for R_1_ in left premotor cortex (cluster spanning left precentral gyrus and left middle frontal gyrus; x = −35, y = 12, z = 34; k = 426 voxels) and for PD in the left postcentral gyrus (x = −66, y = −15, z = 31; k = 242 voxels). For the opposite contrast, significant clusters were found for PD in the left premotor cortex (cluster located in the left superior frontal gyrus; x = −21, y = 1, z = 66; k = 234 voxels), and for R_2_* two clusters in the left cerebellum (x = −12, y = −69, z = −56; k = 258 voxels and x = −26, y = −74, z = −52; k = 375 voxels). At baseline, the training and control groups were comparable across most clusters (Table S4), however, significant between-group differences at S1 were observed for PD in the postcentral gyrus cluster (*p* = .021, Cohen’s d = −0.59) and R_2_* in one cerebellar cluster (*p* = .001, Cohen’s d = 0.83; Table S4). None of the changes in qMRI parameters within the significant clusters were significantly correlated with Session 2 performance on the trained chopstick grasping task (all *p*s > .05; Table S5).

## Discussion

Animal research is widely used to study mechanisms that are assumed to also underlie brain plasticity in humans. However, attempts to directly compare plasticity across species remain rare (Hille et al., 2024). To overcome this impasse, we developed a human analogue of the widely used single-pellet reaching task in rodents (e.g., Xu et al., 2009), preserving its core structural features. We found similarly shaped learning trajectories in both species, indicating that the aligned tasks captured comparable aspects of motor skill acquisition. We applied longitudinal sMRI in both mice and humans to examine training-related brain changes at a shared macroscopic level of analysis. In mice, regional volume in contralateral primary motor and forelimb somatosensory cortex decreased in trained relative to control mice during early learning and greater volume reduction in the contralateral primary motor cortex was associated with better performance. In humans, VBM-based analyses revealed no changes in estimated gray matter volume, whereas analyses of qMRI maps showed increases in myelin-sensitive MT_sat_ and R_1_ estimates in the contralateral premotor and prefrontal cortices during early learning. Together, these findings establish a human analogue of a standard rodent motor learning task and show that such aligned cross-species paradigms, combined with longitudinal MRI, can help facilitate the study of experience-dependent plasticity across species.

### Establishing a cross-species model of motor skill acquisition

#### Designing a human analogue of the rodent reaching task

Our aim was to develop a task for human participants that preserved the core features of the well-established rodent single-pellet reaching task. Several parameters of the task design had to be considered to achieve a similar level of required effort, complexity, motivation, and neural engagement across species. Therefore, the human task was designed as a unimanual dexterity task moving appetitive food items with a reach-grasp-retrieve sequence. It consisted of daily practice divided into discrete trials that were matched in number to the rodent reaching paradigm. However, we made several changes when adapting the rodent task for use in humans. To adjust difficulty for human participants and still retain ecological validity, we introduced chopsticks as a tool. Also, while both tasks used food items, humans were not food-restricted and were only allowed to consume the M&Ms after finishing the task; hence, reward was immediate in mice but delayed in humans.

#### Behavioral validation of the human task

The human behavioral results demonstrated robust motor skill learning following training with the newly developed chopstick grasping task. Participants who underwent daily chopstick grasping training showed pronounced improvements in performance on the trained 30 M&Ms task which exceeded those made by the control group due to repeated task exposure. Importantly, improvements were not restricted to the specific task practiced during training but showed near transfer to related chopstick grasping tasks involving different and more challenging object shapes and use of the non-dominant hand. In contrast, we found no evidence for far transfer to more general measures of fine motor dexterity. This suggests that training produced a specific motor skill extending to closely related grasping demands rather than a broad improvement in manual dexterity.

Notably, the shape of the learning trajectory was similar for rodents and humans, with improvements most pronounced early in training and a decline in session-to-session gain toward the final sessions as performance approached an asymptote in both species. Although both species underwent daily training with the same number of grasping trials per session, the overall training duration differed markedly, spanning 70 days in humans compared to 11 days in mice. Thus, similarly shaped curves do not necessarily imply comparable time bases (Dayan & Cohen, 2011). These findings establish the newly developed chopstick grasping paradigm as a suitable human model of motor skill acquisition, producing robust and measurable behavioral learning while retaining structural correspondence to the single-pellet reaching task used in mice.

### Structural adaptations during experience-dependent plasticity

#### Designing imaging paradigms across species

In both rodents and human participants, we aimed to investigate macroscopic structural brain changes during learning, for which the selection of MRI time points is critical. Some structural changes may emerge early and then renormalize later during training (Lindenberger & Lövdén, 2019; Xu et al., 2009). We attempted to match imaging timepoints to the same learning stage in both species. To cover early plastic changes, we therefore had two training sessions (plus one session to verify paw preference) in mice and six training sessions in human participants between the baseline and the first MRI acquisition after training started. To cover the rest of the learning period, mice were scanned at three-day intervals, while human participants were scanned weekly during the first month and approximately fortnightly thereafter.

Several differences in the imaging procedure between mice and humans were unavoidable in this study due to species-specific constraints. While human participants can be instructed to remain still in the MRI scanner, mice have to be anesthetized to perform longitudinal scans, which may influence physiology, neural activity, and potentially MRI-derived measures (Bajwa et al., 2019; Berkowitz et al., 2020). In addition, handling, repeated scanning, and task training could introduce stress-related effects that potentially act against and are difficult to disentangle from training-related plasticity (Hildebrandt et al., 2008; Kassem et al., 2013). To control for these factors, the mice in the control cohort were also handled and placed into the grasping box at the same frequency and duration as the animals undergoing forelimb training. We additionally introduced a cohort of mice that underwent training but was scanned only at the baseline and final imaging timepoint (anesthesia control group), reducing the number of repeated scanning sessions under anesthesia. In trained animals the total brain volume decreased by 0.7 % over the full interval with two MRI scans, against 1.0 % with four scans, suggesting that part of the global decrease scales with the number of anesthesia sessions; the remainder, present even with two scans, may reflect the food restriction that all groups received.

#### Training-related structural change in mice

While we predicted an expansion-renormalization trajectory in the mouse sensorimotor cortex, we found no significant Time × Group interaction in either analyzed region during the full training period. Contrary to the predicted expansion, the analysis of the early learning interval, revealed a more negative volume change in trained than in control mice in both regions. The increase in gray matter volume reported by Badea et al. (2019) after 25 days of single-pellet reaching training does not necessarily contradict this finding: with only two MRI measurements spanning the entire training period, their design could not resolve early changes. Mediavilla et al. (2022) attributed the decrease in gray matter volume in motor and somatosensory cortex to increased intracortical myelin rather than tissue loss, supported by unchanged histological cortical thickness. Their data, however, cannot exclude gray matter tissue loss, as the two explanations are not mutually exclusive and a histological thickness measure might not detect a change of a few percent. The pipeline used here does not classify voxels within the cortical region of interest as gray or white matter, so a tissue reclassification of the kind suggested by Mediavilla et al. (2022) cannot directly explain the relative volume decrease we observed. Our results therefore point towards a reduction in regional tissue volume in trained mice relative to controls, most plausibly driven by microstructural remodeling rather than by net loss of cells.

That greater volume reduction in primary motor cortex correlated with better reaching performance is consistent with this remodeling being linked to early learning. Several cellular processes may be involved during this time window: myelin sheaths retracting during the first days of motor training, with new myelin appearing only days to weeks later (Bacmeister et al., 2020, 2022); dendritic spines being formed and eliminated rapidly in contralateral motor cortex over the same period (Xu et al., 2009; Yang et al., 2009); and glial compartments remodeling during learning in ways that have been linked to MRI-detectable structural change (Sagi et al., 2012). What each of these processes contributes to a macroscopic volume measure remains unresolved.

#### Training-related structural brain changes in humans

Contrary to our expectations, VBM-based analyses did not reveal grey matter volume changes with training in humans, unlike those reported in previous longitudinal studies of motor skill learning (Draganski et al., 2004; Driemeyer et al., 2008; Gryga et al., 2012; Taubert et al., 2010; Wenger, Kühn, et al., 2017). However, these studies differed considerably in the type of task, training intensity, intervention duration and MRI sampling, making it difficult to determine the temporal window in which training-related gray matter volume changes are most likely to be detectable (Lövdén et al., 2013). Importantly, gray matter volume changes have also been reported within a single session (Kodama et al., 2018; Olivo et al., 2022; Taubert et al., 2016), so our first post-training scan, after six practice sessions, may have missed very early changes, including a potential initial expansion and renormalization occurring within the first week of training (Lindenberger & Lövdén, 2019; Lövdén et al., 2020; Wenger, Kühn, et al., 2017). Furthermore, concurrent cellular processes such as synaptic remodeling and myelin-related plasticity (Xiao et al., 2016; Xu et al., 2009) could change tissue composition without a net change in gray matter volume.

Consistent with the possibility that motor learning induces changes in tissue composition without a detectable change in gray matter volume, MT_sat_ and R_1_ increased in the premotor cortex, and MT_sat_ increased in the prefrontal cortex between session 1 and session 2 in the training group. Both measures are sensitive to myelin, MT_sat_ through magnetization transfer between free water and the macromolecular pool, to which myelin contributes substantially, and R_1_ through the influence of macromolecules and lipids on the local relaxation environment of tissue water (Helms et al., 2008; Henkelman et al., 2001; Lutti et al., 2014; Turati et al., 2015; Weiskopf et al., 2013, 2021). To our knowledge, only two previous longitudinal studies of motor skill learning in humans have similarly acquired multiple qMRI parameter maps (Aye et al., 2025; Azzarito et al., 2023). Both reported changes in myelin-sensitive measures across different brain regions, with Azzarito et al. (2023) showing decreases followed by increases and Aye et al. (2025) finding decreased MT_sat_ but increased R_1_ from pre-to post-training. Differences in task, brain regions, and temporal sampling limit direct comparison, with the absence of intermediate measurements in Aye et al. (2025) potentially obscuring earlier or transient changes during learning. Nevertheless, together with our findings, these studies indicate that motor skill training is associated with dynamic changes in myelin-sensitive qMRI measures, potentially reflecting changes in intracortical myelin, although their temporal trajectories and spatial specificity remain to be consistently established.

Although our data do not provide direct information on the underlying cellular mechanisms, oligodendrocyte plasticity is a plausible substrate for the observed increases in MT_sat_ and R_1_. In rodents, new oligodendrocytes appeared in the motor cortex within hours of exposure to a complex wheel-running task, and blocking their formation impaired learning (McKenzie et al., 2014; Xiao et al., 2016), suggesting an important contribution of oligodendrocyte plasticity to early motor learning. Beyond these increases in myelin-sensitive measures, we also observed training-related decreases in R_2_* in the cerebellum. As R_2_* is sensitive to tissue iron (Ghadery et al., 2015; Langkammer et al., 2010), the observed decrease may reflect changes in iron content or distribution accompanying increased metabolic and cellular demands during motor learning, including synaptogenesis and oligodendrocyte plasticity (Azzarito et al., 2023; Callaghan et al., 2014; Carlson et al., 2007; Fretham et al., 2011; Tran et al., 2015).

Together, the changes across qMRI metrics suggest that motor learning may be accompanied by multiple forms of tissue remodeling that are not necessarily reflected in macroscopic gray matter volume changes.

#### Cross-species interpretation of early structural change

Both species showed evidence for early structural changes during training in contralateral sensorimotor regions, though observed with different MRI measures: a regional volume decrease in trained mice relative to controls and an increase in myelin-sensitive MT_sat_ and R_1_ in humans. For the one measure obtained in both species, volume, the results differed: mice showed an early relative decrease while humans showed no significant change. Faced with a similar discrepance, Mediavilla et al. (2022) speculated whether this reflects differences in task, field strength, or MRI sequence. Given that the tasks were rather similar, potential explanations are more likely to refer to the nature and timing of measurement, particularly a later first post-training scan in humans. At the same time, it is worth noting that the volume decrease in mice and the MT_sat_ and R_1_ increase in humans need not be conflicting: the mouse measure reflects regional volume, whereas the human qMRI measures capture tissue composition. The human findings point to myelin-associated remodeling as a candidate mechanism, and intracortical myelin remodeling could also contribute to the relative volume decrease in mice. Unfortunately, the rodent part of this study can neither confirm nor exclude this possibility. Nor does the rodent literature resolve the question: using the same task, Mediavilla et al. (2022) reported a transient increase in intracortical myelin during learning in forelimb somatosensory cortex, whereas Bacmeister et al. (2020, 2022) observed early sheath retraction in motor cortex, a discrepancy that Mediavilla et al. themselves note and that is further complicated by differences in methods and region.

### Limitations

Task demands were not fully matched across species. The principal departure from rodent grasping was the introduction of chopsticks, which was necessary to reach comparable difficulty levels, but could engage partly different brain circuits. Moreover, mice were food-restricted and consumed each millet seed immediately after grasping, whereas human participants were not food-restricted and were only allowed to consume the sweets after task completion. Hence, although seeing each M&M may have served as a reward-predictive cue, motivational state and reinforcement structure likely differed between species, potentially influencing task engagement, motor learning, and associated neural plasticity through differences in reward-related processing, including dopaminergic signaling (Abe et al., 2011; Hosp et al., 2011; Molina-Luna et al., 2009; Wickens et al., 2003; Wise, 2004). Additionally, the session criteria differed: Human sessions ended after 30 successful transfers, mouse sessions after 30 attempts, unless the predefined time limit was reached first. Humans therefore made more grasping movements than mice, particularly early in training. The criteria also led to different outcome measures, with the completion time in humans capturing both speed and errors, whereas success rate in mice captured accuracy only, because completion time could be confounded by motivational state in mice. The learning curves can still be described as similar in shape, supported by comparable rate parameters from the exponential models.

The imaging was less closely matched between species than the behavior. MRI timepoints were chosen to match the learning stage regarding the full training duration, leading to a different training dose in the early interval: six practice sessions in humans against two in mice. It therefore remains possible that structural changes occurring before the first post-training scan, including an initial expansion and subsequent renormalization, were not captured by the present sampling schedule in humans. Further, in mice, the requirement for repeated anesthesia, and possibly the food restriction common to all groups, might have contributed to the global volume decrease, as indicated by the anesthesia control group. And finally, MRI acquisition protocols differed between species, reflecting differences in sequence availability and constraints on acquisition time in anaesthetized animals. In mice, structural images were acquired using a 3D gradient-echo (FLASH) sequence, whereas the imaging protocol for humans included T_1_-weighted structural imaging and quantitative multiparameter mapping. Additionally, morphometric measures were derived differently in the two species. Mouse regional volumes were obtained by multi-atlas label fusion segmentation, and the analysis was restricted to two ROIs, whereas in human participants, the analysis was performed voxel-wise across the whole brain using VBM. These methodological differences should be considered when interpreting apparent similarities or differences across species.

### Future directions

Several improvements to the present design follow from these limitations. The clearest is the acquisition of myelin-sensitive sequences in mice, potentially as part of a qMRI protocol, which would align measures more closely between the species (Rahman et al., 2023). In addition, mouse structural scans should be optimized for VBM, so that the same analysis approach of gray matter volume change can be applied in both species. Denser MRI sampling early in training in the human cohort could improve temporal alignment with the early training window sampled in mice and capture potential rapid structural changes occurring before the first post-training scan (Olivo et al., 2022; Xu et al., 2009), including a transient initial gray matter volume expansion (Lindenberger & Lövdén, 2019; Lövdén et al., 2020).

To further align behavioral procedures across species, modifications to the motivational state, reward structure, training dose, and outcome measure should be considered. Future studies could seek to match motivational states by reducing or removing chronic food restriction in animals, which could possibly also reduce the total brain volume decrease observed in mice. Indeed, in a separate pilot test (data not shown), mice that had fasted for up to six hours before each session were still able to learn the reaching task. On the human side, alignment could be improved by trying to standardize and potentially enhance human participants’ motivation to obtain food rewards. Possible approaches include scheduling training at a consistent time relative to meals, implementing controlled meal schedules or mild food restriction and assessing subjective appetite before and after the task, as well as trying to account for individual differences in preference for the food reward (e.g., M&Ms). Additionally, reward structure could be aligned more closely by allowing human participants to consume the sweets after individual successful trials or short trial blocks, while care needs to be taken to avoid the consumption confounding completion time as an outcome. To match the training dose, it would be possible to either allow humans to only perform 30 attempts, like mice, or to use 30 successful retrievals as the end point in the rodent experiments. However, sessions in mice ending after 30 successes would likely become very long early in training, making task adherence more dependent on motivational state. Success rate could be a common outcome measure in both species but would require a careful definition of what constitutes a grasping attempt in the human cohort. Finally, video-based motion capture would allow one to study the kinematics of reach-to-grasp movements and their change with training, providing an additional behavioral measure that is comparable across species (Mathis & Mathis, 2020).

As proposed in greater detail by Hille et al. (2024), the major goal of experimental paradigms matched across species is to advance the mechanistic interpretability of brain changes observable in humans. Macroscopic MRI changes are the net outcome of cellular processes that can be measured directly in mice. Candidate cellular mechanisms for the present task include dendritic spine formation and elimination (Fu et al., 2012; Xu et al., 2009), oligodendrogenesis and myelin remodeling (Bacmeister et al., 2020; McKenzie et al., 2014; Xiao et al., 2016), and astrocyte plasticity (Kleim et al., 2007; Padmashri et al., 2015). Longitudinal two-photon imaging of these processes has already been established for dendritic spines (Xu et al., 2009) and oligodendrocytes (Bacmeister et al., 2020) during learning. Combined with structural MRI, in vivo microscopy could help link the macroscopic measure to its cellular substrate in the same brain, as has been done for cell density and nucleus volume (Asan et al., 2021). Astrocytes are also a promising target: an experience-dependent increase in gray matter volume in the visual system has been attributed mainly to astrocyte swelling (Schmidt et al., 2021), yet, to our knowledge, astrocyte morphology has not been followed longitudinally in vivo during motor learning. Additionally, candidate processes, including oligodendrogenesis or spine formation, can be experimentally manipulated rather than only observed (Joag et al., 2026; Xiao et al., 2016), to establish whether a macroscopic volume change depends on a particular cellular process or merely accompanies it. None of this is possible in humans. With behavior and imaging aligned between mice and humans, a mechanism established in mice becomes a candidate explanation for the human signal rather than a mere analogy to it.

Future cross-species studies would also benefit from MRI measures beyond volume. Frameworks that seek to integrate microscopic and macroscopic findings, such as the expansion– exploration–selection–refinement (EESR) model (Hille et al., 2024; Lindenberger & Lövdén, 2019; Lövdén et al., 2020), propose a temporal sequence of plastic changes, and several of the processes involved can be measured in both species. In mice, it has been shown that during motor learning the population of movement-related neurons in the mouse motor cortex first expands and then refines into a smaller and stable ensemble with reproducible activity (Peters et al., 2014). In humans, repeated task fMRI could test whether task-related activation is heightened early in learning and precedes structural brain changes, as predicted by the EESR model. Furthermore, representational similarity analysis (Kriegeskorte et al., 2008) could be used to examine whether task-relevant multivoxel activation patterns become more stable over the course of motor training. Previous findings in mice indicate that reduced cortical inhibitory tone may facilitate plasticity induction in the motor cortex during motor skill acquisition (Chen et al., 2015). In humans, functional magnetic resonance spectroscopy could track glutamate and γ-aminobutyric acid (GABA) concentrations in task-relevant motor regions, providing information about neurochemical changes potentially related to a shift in excitation–inhibition balance that may precede structural changes (Floyer-Lea et al., 2006; Kolasinski et al., 2019; Stanley & Raz, 2018). Measured at matched time points in both species, together with the cellular and myelin-sensitive MRI measures outlined above, these markers could help determine the relative timing of plastic changes during motor skill acquisition, and thereby test, and potentially refine, the sequence of changes proposed by the EESR model.

## Conclusion

We designed a human analogue of the rodent single-pellet reaching task and found that learning in both species followed similarly shaped trajectories, although on different time scales. Combined with longitudinal MRI, the matched paradigm revealed early structural changes in contralateral sensorimotor regions in both species: a regional volume decrease in trained relative to control mice, with larger decreases in primary motor cortex associated with better reaching performance in trained mice, as well as changes in myelin-sensitive tissue properties in trained humans. Whether these findings reflect a shared biological process cannot be resolved with the measures available in this study. The contribution of this work is the matched behavioral paradigm: a task that both species learn comparably is the precondition for asking whether the underlying biology is shared and allows the observations and manipulations possible in mice to be leveraged to interpret human brain plasticity more directly. An important next step in this line of work will be to incorporate longitudinal two-photon microscopy into the mouse protocol. As proposed elsewhere (Hille et al., 2024), this would allow cellular changes observed during learning to be related directly to MRI-derived structural changes in the same animals and, in turn, to corresponding structural changes observed in humans.

## Material and Methods

### Participants

#### Mice

A total of 51 male C57BL/6N mice (selected to align the sex composition with the all-male human cohort), aged 8 weeks at the start of the experiment, were used: 19 animals in the training group, 17 in the control group and 15 in the anesthesia control group. Two animals in the training group and one animal in the control group were eliminated from MRI analysis because it was not possible to acquire four artifact-free scans, but the trained mice were retained for behavior analysis. Additionally, two of the animals in the anesthesia control group failed to learn the task and were excluded from all further analysis. Animals were single-housed in IVC II cages and kept at 22 ± 1.5 °C and 55 ± 5% relative humidity under an inverted light/dark cycle with ad libitum access to water. To motivate reaching, all animals, including controls, were food-restricted to a target weight of about 85–90% of their free-feeding baseline body weight from day −10, 11 days before the first training session. They usually received 2–3 g of chow per day, were weighed daily, and welfare was monitored with a score sheet. All animal handling and procedures were conducted in accordance with the ethical guidelines established by the Max Planck Society and protocols approved by the Regierung von Oberbayern (Beratende Ethikkomission nach §15 Tierschutzgesetz).

#### Humans

Seventy-five male participants aged between 18 and 35 years (*M* = 24.4, *SD* = 4.74) were recruited through flyers, via the online platform Kleinanzeigen (https://www.kleinanzeigen.de/), a web platform for study recruitment (https://www.probando.io/), and a participant database developed for the Max Planck Society (Bengfort et al., 2022). All participants were right-handed, had normal or corrected-to-normal vision, no diagnosis of psychological or neurological diseases, no contraindications for MRI studies, and little to no experience with chopstick use. The sample included only male participants to minimize the potential hormonal influence of the menstrual cycle on brain structure and function during the intervention (Lisofsky et al., 2015; Pletzer et al., 2010). Prior to the start of the study, participants were randomly assigned to either the chopstick training group or the reading control group. Four participants discontinued the study after the first or during the second appointment and were therefore excluded from data analysis. Seventy-one participants remained for analysis: 36 participants in the chopstick training group (*M* age = 24.6 years, *SD* = 4.88) and 35 participants in the reading control group (*M* age = 24.1 years, *SD* = 4.68). The exact sample sizes used in each analysis are reported in the relevant sections below, as not all participants completed the full training and all in–person sessions. The study was approved by the local Psychological Ethical Committee at the Center for Psychosocial Medicine at University Medical Center Hamburg-Eppendorf in Hamburg, Germany (LPEK-0273 and LPEK-0380). Participants provided written informed consent and received financial compensation for their participation in the study.

### Experimental Design

#### Mice

Mice were randomly allocated to three groups (training, control and anesthesia control). The training group as well as the anesthesia control group were trained on the single-pellet reaching task. The control group received identical handling but did not undergo reach training. The training and control groups were each scanned four times (S1–S4: day −3, 3, 7, 11), while the anesthesia control group was only scanned twice (day −3 and 11). The experiments were carried out in two batches, each including mice of all three groups.

Days are numbered relative to the first training session (day 1). First, the paw preference was assessed on day −18, followed by the headbar implantation on day −17. Food restriction started on day −10, after 7 days of recovery. A second paw preference test was done on day 0. Training took place on days 1, 2, 4, 5, 6, 8, 9 and 10.

##### Headbar implantation

On day −17, all animals were implanted with an MRI-compatible plastic headbar (25 × 15 × 1 mm with a central opening; custom-made by the workshop of the Max Planck Institute of Psychiatry). The headbar allowed the receive coil to be clipped in the same position at every MRI session. Mice were anesthetized with fentanyl, midazolam and medetomidine (0.05, 5, and 0.5 mg/kg, i.p., respectively), received carprofen (5 mg/kg, s.c.) for analgesia and 1 ml of sterile saline subcutaneously for fluid replacement. They were fixed in a stereotaxic frame; their eyes were protected with Bepanthen eye and nose ointment (Bepanthen® Bayer AG, Leverkusen, Germany), and their body temperature was kept at 37.5 °C with a feedback-controlled heating pad. After incision of the scalp, lidocaine was applied for local anesthesia, and subsequently the skull exposed. The headbar was fixed with dental cement (Paladur), which was applied with a small spatula to avoid air bubbles. Anesthesia was antagonized with naloxone, flumazenil, and atipamezole (1.2, 0.5 and 2.5 mg/kg, i.p., respectively). Carprofen injection (5 mg/kg, s.c.) was repeated on the evening of surgery and the next morning, followed by carprofen in the drinking water (20 mg/kg) for a further 24 h. No craniotomy was performed. Animals recovered for seven days before food restriction began on day −10. In some cases the plastic headbar detached from the dental cement and was re-glued with superglue during the anesthesia for the following MRI acquisition.

##### Single-pellet reaching task

Mice were trained on the single-pellet reaching task adapted from Xu et al. (2009). They were placed in a transparent acrylic glass box (approximately 15 × 8.5 cm) with three vertical slits; a central slit on one side for the paw preference test and two lateral slits on the other side for the training sessions. For the paw preference test, which also served as habituation to the box, millet seeds were presented on a spoon in front of the central slit. Paw preference was determined when mice had made at least 15 reach attempts and showed > 50% limb preference. This preference test was done twice, once before the headbar implantation, on day −18, and once immediately before the start of training, on day 0. Paw preference was as follows: Training group 10 right, 7 left preferring; anesthesia control group 10 right, 3 left preferring; control group 7 right, 5 left preferring, and 4 without detectable preference — either ambiguous or not enough reaching attempts during the test. During the training sessions, a single millet seed was presented on a small spoon in front of the lateral slit, corresponding to the paw preference of the animal. The seed was precisely positioned in front of the slit with a custom-built device. A reach was scored as successful when the mouse grasped the seed with the preferred paw, brought it through the slit and ate it without dropping it. One session was run per day, with a maximum length of 20 min or comprising 30 reach attempts. Mice were trained daily, except on MRI days, for a total of eight sessions over 11 days (training days 1, 2, 4, 5, 6, 8, 9, and 10). Control mice received identical handling and spent 20 min per day in the box, with 30 millet seeds placed inside.

#### Humans

The study followed a longitudinal within-subject design over 10 weeks (70 days). Each participant completed an online questionnaire and was invited to seven in-person sessions at the Max Planck Institute for Human Development (on days 1, 8, 15, 22, 38, 54, and 70; Figure 2A). At every in-person session, all participants underwent MRI measurements. Behavioral assessments of chopstick grasping performance and general fine motor skills were administered only during the first and last sessions. Throughout the entire study period, participants engaged in a daily home-based activity, performing either a standardized chopstick grasping task or a matched-duration reading task (see below for details).

##### Online questionnaire

Prior to the first in-person appointment, participants completed an online questionnaire that collected demographic information (age, gender, education), information on handedness, self-reported experience with chopstick use, and experience with other fine motor skills (e.g., playing a musical instrument or 10-finger typing). Additionally, the questionnaire included the behavioral inhibition system and behavioral approach system (BIS/BAS) scales (Strobel et al., 2001), the chocolate-specific versions of the Food Cravings Questionnaires (Meule & Hormes, 2015), and the 10-item version of the Achievement Motives Scale (Lang & Fries, 2006).

##### MRI measurements

All participants were invited for seven in-person sessions at the institute on days 1, 8, 15, 22, 38, 54, and 70 for MRI data acquisition. The MR measurement protocol included a T1-weighted structural whole brain scan, single-voxel spectroscopy of the primary motor cortex, a high-resolution proton density sequence for hippocampal subfield measurements, task-based and resting state fMRI, a multiparameter mapping (MPM) protocol and diffusion-weighted imaging (DWI). The present study reports analyses of a subset of these acquisitions (see MRI Data Acquisition below for the corresponding sequence parameters). To minimize potential short–term training-related effects on brain activation or structure, participants were instructed not to perform their daily training before the scanning sessions.

In the first session, we administered a finger-tapping task inside the scanner. Participants were instructed to tap the thumb and index finger of the indicated hand against each other at a rate of approximately 1 Hz. Blocks requiring movements of the left or right hand were cued by the letters “L” and “R” presented on a screen. This task served as a functional localizer to map the cortical representations of the left and right hands (Gelnar et al., 1999; Wenger, Kühn, et al., 2017).

In addition, participants performed an fMRI task designed to probe the neural correlates of the trained chopstick grasping behavior. This task was administered to all participants during the first and last sessions, and in the chopstick training group in all intermediate sessions as well. It served to identify brain regions functionally involved in the execution of the trained grasping behavior over the course of training and was used solely for functional localization, without acquisition of a behavioral outcome measure. For the in-scanner chopstick task, participants lay supine with a custom-built wooden apparatus placed on their abdomen. The apparatus consisted of a flat wooden base board and a perpendicular semi-circular vertical panel mounted on it. A wooden rod was attached to the vertical panel parallel to the participant’s body, and a wooden marble with a central hole was mounted on the rod so that it could be translated along its length. Using either a pair of chopsticks in the right hand (chopstick condition) or the right hand without chopsticks (hand condition; using thumb, index, and middle finger), participants were instructed to repeatedly grasp, move, and release the marble along the rod for as long as the corresponding cue was present. The apparatus was dimensioned to allow a comfortable posture while restricting movement primarily to the wrist and fingers, thereby minimizing upper-arm and head motion. The head coil was equipped with two mirrors that allowed participants to view their hand and the apparatus while keeping the head stable. Task conditions were indicated by colored light projected into the scanner bore: blue for the chopstick condition, orange for the hand condition, and green for rest. The first block consisted of 12 chopstick trials interleaved with 6 rest trials (10 s per trial, randomized order), and the second block consisted of 12 hand trials interleaved with 6 rest trials (7 s per trial, randomized order). The chopstick trials were intentionally longer to allow participants sufficient time to re-establish their grip and initiate the movement. A jittered fixation cross (4–11 s) was presented between trials. During rest trials of the chopstick block, participants held the chopsticks passively without putting them down to further reduce movement.

##### Behavioral assessment at the institute

Behavioral assessments were conducted in a quiet testing room following MRI acquisition. At S1 and S7, all participants completed the same set of tasks in the same order. Standardized instructions were read aloud, and task order was fixed across participants, with short breaks provided as needed to minimize fatigue. The behavioral battery therefore provided matched pre- and post-intervention measures across groups.

##### Chopstick grasping tasks

Participants first completed a chopstick grasping task (Figure 1) that was identical to the daily training task performed at home (see “Daily training” below). Using a standard pair of chopsticks in their dominant right hand, they were instructed to grasp and transport 30 M&Ms (chocolate-coated peanuts) from a start location to a target container as quickly and accurately as possible. The time required to transport each set of 10 M&Ms (1–10, 11–20, 21–30) as well as the total time for all 30 M&Ms was recorded, with a maximum allowed duration of 21 minutes. If participants did not complete all 30 transfers within this time, the number of successfully transported M&Ms was noted. Participants in the chopstick training group performed this task at each in-person session after the MR assessment, whereas participants in the control group only performed this task at S1 and S7. All other behavioral tasks described below were administered at S1 and S7 in both groups.

Subsequently, participants performed several additional chopstick grasping tasks. The procedure was analogous to the task described above but with different objects. Participants had to grasp and transport 10 glass marbles with chopsticks in their right hand (maximum duration: 7 min), 30 Maltesers (round chocolate candies) with chopsticks in their right hand, and 10 M&Ms with chopsticks in their left hand (maximum duration: 7 min). Participants also performed a modified version of the Purdue Pegboard Test using chopsticks with their right hand. Instead of placing pins using their fingers, participants used chopsticks to pick up and insert the pins into the board. The outcome measure was the number of pins correctly placed within 60 s. These tasks were designed to probe the transfer of training-related changes in chopstick grasping to different objects and, for M&Ms, to the other hand.

##### General fine motor skills

To assess general manual dexterity participants completed standardized tests at sessions 1 and 7: Purdue Pegboard Test (Tiffin & Asher, 1948): Performed with the right hand, left hand, both hands, and in an assembly condition following the standard administration procedure. The primary outcomes were the total number of correctly placed pins (unimanual condition), the total number of pairs of pins inserted (bimanual condition) and the number of parts assembled (assembly condition) within the allotted time.

Grooved Pegboard Test (Matthews & Klove, 1964): Performed separately with the right and left hand according to standard instructions. Th outcome measure was the time to complete the board.

O’Connor Tweezer Dexterity Test (O’Connor, 1928): Performed with the right hand using tweezers to place small pins into holes. The outcome measure was completion time.

## Daily training

### Chopstick training group

Participants in the chopstick training group performed a standardized, experimenter-developed chopstick grasping task at home on each of the 70 study days. The grasping task was adapted from the single-pellet reaching task in mice in order to have very similar training regimes for both species. For this purpose, participants were provided with a custom setup (wood / acrylic glass apparatus), two glass bowls, a pair of chopsticks, a supply of M&Ms (chocolate-coated peanuts), and a training diary. Each day, participants placed 30 new M&Ms into the left bowl and then sequentially transported them from the left to the right bowl. On each trial, a single M&M was positioned centrally behind a narrow slit with the left hand, and participants used chopsticks in their right hand to grasp and transport the M&M through the narrow slit and then place it into the right bowl. This procedure was repeated until all 30 M&Ms had been transferred. Participants recorded the time required to complete the task and noted it in the training diary as the primary outcome measure. Participants were instructed to wash and thoroughly dry their hands before training to avoid slippage. They were allowed to consume the M&Ms after completing the task. Before each session, participants self-reported sleep quality and training motivation in the training diary and after training, they documented task completion. In order to monitor performance and compliance, participants placed their smartphone on top of the apparatus and recorded a video of each training session, which they uploaded once per day via a secure online link.

### Reading control group

Participants in the control group engaged in a matched-duration reading task instead of chopstick training. They were instructed to read for approximately 10 minutes each day over the 70-day study period, corresponding to the average daily training time of the chopstick training group. All control participants received the same model of e–reader preloaded with seven different books. Participants were free to choose which of the provided books to read on any given day and could determine the order in which they read them across the study period. To monitor compliance, participants completed a brief daily diary in which they reported their reading duration, the book they had read, and additional ratings related to the reading session. Participants were asked to upload a photograph of the completed diary page once per week via a secure online link.

### Behavioral Data Analysis

All analyses were performed in R 4.2.2 (R Core Team, 2022) using RStudio (Posit team, 2026) and the following packages: tidyverse (Wickham et al., 2019), lme4 (Bates et al., 2015), lmerTest (Kuznetsova et al., 2017), emmeans (Lenth, 2022), performance (Lüdecke et al., 2021) and DHARMa (Hartig, 2016).

#### Mice

Behavioral data from the grasping task performed by trained mice (*n* = 32; anesthesia control: *n* = 13) on each training day were analyzed using generalized linear mixed-effects models. For each training day, we used the number of successful reaches out of 10 attempts within each set of consecutive reaches (reaches 1–10, 11–20, 21–30), hereafter referred to as bins and modeled success counts with a binomial family and logit link to appropriately account for the bounded, count-based nature of the outcome. We initially fitted a simple model containing only linear training day effects and then progressively increased complexity comparing nested models using likelihood-ratio tests. The final model included fixed effects of training day (linear and quadratic terms), bin, and their interactions, random intercepts for animals (capturing individual differences in overall performance), and random slopes for training day per animal (capturing individual differences in learning rate). Training day was mean-centered in this final model to reduce collinearity between linear and quadratic terms and their interactions with bin. Model assumptions were assessed via residual diagnostics, including quantile-residual simulation using DHARMa, visual inspection of residual and Q-Q plots, and collinearity diagnostics. We assessed which consecutive training day intervals showed the strongest performance improvements by estimating marginal mean performance at each training day, averaged across bins, and comparing consecutive training days pairwise (e.g., day 2 vs. day 1, day 4 vs. day 2). Within training day, differences across bins were evaluated by estimating marginal means for each bin and conducting pairwise comparisons between successive bins, that is, bin 2 versus bin 1 and bin 3 versus bin 2. Holm correction was used to account for multiple comparisons.

#### Humans

To test whether training was successful and the chopstick training group improved more than the reading control group in the chopstick grasping tasks, we analyzed performance at the first (S1) and last sessions (S7). Analyses were restricted to participants who had completed both pre– and post–training sessions (*N* = 65; chopstick training group: *n* = 33; reading control group: *n* = 32). For each task, we fitted a mixed–effects model with fixed effects of group (chopstick training vs. reading control), session (S1 vs. S7), their interaction, and a random intercept for participants. For the 30 M&Ms and 30 Maltesers tasks, we used linear mixed-effects models with the log-transformed completion time as the dependent variable. For the 10 marbles and 10 M&Ms (left hand) tasks, we analyzed the number of successful grasps out of up to 10 items with beta-binomial mixed-effects models. For the modified version of the Purdue Pegboard test using chopsticks, we used a zero–inflated Poisson generalized linear mixed–effects model. We also tested for group differences in the more general fine motor dexterity tasks. We used linear mixed-effects models for the Purdue Pegboard test, Grooved Pegboard test and the O’Connor Tweezer dexterity test. For the latter two models, the outcome variable (total completion time) was log–transformed. Model assumptions were checked using residual plots, Q–Q plots, and diagnostic tools from the performance and DHARMa packages.

In addition to these pre–post group comparisons, we analyzed session-wise learning curves in the chopstick training group (*n* = 36 participants) to characterize how performance changed over the course of training. We analyzed behavioral data from the chopstick grasping task performed at each in–person session using linear mixed–effects models. For each session, we used the recorded times required to transport each set of 10 M&Ms (1–10, 11–20, 21–30) and analyzed these times on the log–transformed scale to satisfy model assumptions of normality and homoscedasticity of residuals. We started with a simple mixed model containing only linear session effects and gradually increased complexity, comparing nested models with likelihood–ratio tests. The final model included fixed effects of session (linear and quadratic terms), bin, and their interactions, random intercepts for participants (capturing individual differences in overall speed), and random slopes for session per participant (capturing individual differences in learning rate). Session number was mean-centered in this final model to reduce collinearity between linear and quadratic terms and their interactions with bin. Model assumptions were assessed via residual diagnostics, including quantile-residual simulation (DHARMa), visual inspection of residual and Q–Q plots, and collinearity diagnostics. To identify between which consecutive training sessions performance improved most strongly, we estimated marginal mean performance at each session (averaged across bins) and performed pairwise comparisons between consecutive sessions (e.g., S2 vs. S1, S3 vs. S2). To assess performance differences between bins within each session, estimated marginal means were calculated for each bin, followed by pairwise comparisons between successive bins (bin 2 vs. bin 1; bin 3 vs. bin 2). We applied Holm correction for multiple comparisons.

#### Characterization of Learning Curves Across Species

To characterize the shape and rate of the overall learning trajectory across sessions in each species, separate mixed-effects models were fitted to session-level overall performance, without the within-session bin factor but otherwise identical to the models described above. Population-level performance estimates across sessions from these models were then fitted separately for each species with an exponential function including three parameters representing asymptotic performance, the initial deviation from asymptotic performance, and the learning rate k. This function provided a compact description of the population-level learning trajectories and a single parameter summarizing the rate of learning.

### MRI Data Acquisition

#### Mice

Mice were anesthetized with isoflurane (Isofluran CP®, CP-Pharma®, Burgdorf, Germany): Sedation was initiated with 2.5% isoflurane in 1.5 l/min air and maintained during the MR scanning in the range 1.5%–2.5%. The range of breaths per minute was kept between 80 and 120, and depth of anesthesia was adjusted to match this range. Body temperature was kept between 36 and 37.5°C, using a heating pad (water bath, Haake S 5P, Thermo Fisher Scientific, Waltham, United States).

Animals were placed in a prone position on an animal bed with a pressure pillow to monitor respiration and rectal temperature probe. The head was fixed using the headbars. Ophthalmic ointment (Bepanthen® Bayer AG, Leverkusen, Germany) was applied on their eyes to prevent damage due to drying during scanning. After the successful acquisition, anesthesia was discontinued and the mouse was returned to its home cage. Recovery of the animals was monitored. Total time of anesthesia per scanning session was 65–90 min per mouse.

Anatomical images were acquired on a Bruker BioSpec 94/20 MRI scanner (9.4 T) using ParaVision software (6.0.1, Bruker, Ettlingen, Germany). Data were collected using a whole-body coil for RF transmission and a room temperature surface coil (diameter 10 mm) for signal detection. The surface coil could be attached to the headbar which guaranteed the same positioning of the coil relative to the skull for all individual scanning sessions. After scanner preadjustments, a low-resolution multislice localizer scan was run covering all three major slice orientations. Shimming was performed automatically by optimization of the free induction decay (FID) of the global signal. Finally, anatomical scans were run, using a fat suppressed 3D FLASH sequence with repetition time (TR) = 20 ms, echo time (TE) = 5.86 ms, flip angle 10°, matrix size 200 ×140 × 100, field of view 20 × 14 × 10 mm^3^, resulting in an isotropic voxel resolution of 0.1 mm. RF-spoiling was used. Four averages were acquired per run, and the 3D experiment was run 3 times.

#### Humans

All MR images were obtained from participants laying in a supine position, using a CE-certified 3-Tesla TIM Trio MRI scanner (Siemens Medical Systems, Erlangen, Germany) equipped with a 32-channel head coil.

Structural images were acquired using a three-dimensional T1-weighted magnetization prepared gradient-echo sequence (MPRAGE). The following parameters were used: TR = 2500 ms, TE = 4.77 ms, inversion time (TI) = 1100 ms, flip angle = 7°, bandwidth = 140 Hz/pixel, acquisition matrix = 256 mm × 256 mm × 192 mm, isometric voxel size = 1 mm^3^, acquisition time = 9.20 min.

Whole-brain functional images were collected with a T2*-weighted single shot echo-planar imaging (EPI) sequence sensitive to BOLD contrast with the following parameters: TR = 2000 ms, TE = 30 ms, flip angle = 80°, distance factor = 20%, isometric voxel size = 3 mm^3^, FOV = 216 × 216 × 129 mm^3^, using GRAPPA acceleration factor 2 over an acquisition time of 10.08 min. Thirty-six axial slices were acquired in an interleaved fashion, aligned to the genu splenium of the corpus callosum.

A fast 1.6-mm isotropic quantitative multi-parametric mapping (MPM) protocol (Cooper et al., 2020) was used to acquire the following whole-brain MPM maps: proton density (PD), magnetization transfer saturation (MT_sat_), longitudinal relaxation rate (R_1_), and transverse relaxation rate (R_2_*). The parameters used for these sequences are described in detail in Cooper et al. (2020).

### MRI Preprocessing and Statistical Analyses

#### Mice

For each mouse and session, the three repeated acquisitions were realigned and averaged in SPM12. Images of mice that performed the task with the left forelimb were then mirrored across the midsagittal plane, so that the hemisphere contralateral to the trained forelimb corresponded to the left hemisphere in all images. The registration and segmentation of the MRI scans were performed using the pydpiper toolkit (Friedel et al., 2014). First, all images were rigidly aligned to a single study scan. Successive affine and nonlinear alignments registered all scans to a common space and created a study-specific template (Avants et al., 2008). Segmentation of the MRI scans was performed using MAGeT (Chakravarty et al., 2013), with the DSURQE atlas (Dorr et al., 2008; Qiu et al., 2018; Richards et al., 2011; Steadman et al., 2014; Ullmann et al., 2013) registered to the study-specific template serving as the input atlas. The final label of each voxel was determined by majority vote across the candidate segmentations derived from the atlas and the template images. Volumes of the different ROIs were calculated by counting the number of voxels assigned to each label and multiplying by voxel volume. The quality of the segmentation was visually inspected for every scan. In one trained animal, rigid registration of the final scan (day 11) failed, so that atlas labels were placed on the wrong tissue. The scan was excluded from all analyses; the animal’s three other scans were retained. No other scan showed a registration failure.

Analyses were performed in MATLAB R2023b (with Statistics and Machine Learning Toolbox). In this section, the scans on days −3, 3, 7, and 11 are referred to as sessions S1–S4. Two regions of interest were planned, primary motor cortex and forelimb somatosensory cortex, each contralateral to the trained paw (respectively the preferred paw in control mice). Because images of left-pawed mice were mirrored before registration (see Preprocessing), the contralateral hemisphere was the left one in every animal with a paw preference. Control mice without a detectable paw preference were not mirrored, so the left hemisphere was analyzed in every animal (DSURQE labels 269 and 283). The analysis included 17 trained and 16 control mice that were scanned four times each (with exclusion of one scan at S4 in one trained mouse, see Preprocessing), and 13 trained mice scanned only twice (anesthesia control, used for analyses of total brain volume and behavior only).

Total brain volume (the sum of all 336 labeled regions) was compared between groups at baseline with a Student’s t test and across scans with a linear mixed-effects model (fitlme, restricted maximum likelihood) with fixed effects of group (trained, control), time (scans S1–S4, categorical) and their interaction and a random intercept for each animal. Factors were effect-coded (sum-to-zero), so that the main effect of time reflects the change averaged over both groups. Fixed effects were tested with F and t tests using Satterthwaite-approximated degrees of freedom. The Time × Group interaction across all four scans tested for any difference in trajectory between groups. The S1 to S4 change in total brain volume (in percent of each animal’s scan S1 volume) of trained mice scanned four times (*n* = 16) and twice (*n* = 13, anesthesia control group) was compared using Welch’s t test. Group changes from scan S1 to S4 are given in percent of the group’s scan S1 mean, computed on animals with both scans.

Regional volumes were compared between groups using the same mixed model, with total brain volume included as an additional time-varying covariate, and the Time × Group interaction across all four scans was tested. Because structural changes were expected to be largest early in learning (Mediavilla et al., 2022; Xu et al., 2009), the change from baseline (S1) to the first post-training scan (S2), covering the early learning period, was tested as the group × time coefficient for scan S2 in the same model fitted with reference coding (reference levels: control, scan S1), that is, the difference between groups in the S1 to S2 change. For each type of test (omnibus interaction, S1 to S2 contrast and brain–behavior correlation), *p* values were corrected across the two regions with the Holm method; confidence intervals are unadjusted 95% intervals. Each group’s own S1 to S2 change was estimated from the same model (control: scan S2 coefficient; trained: sum of the scan S2 and group × scan S2 coefficients) and tested against zero, with Holm correction across these four tests (two groups, two regions). Regional percent changes are model estimates relative to the mean baseline volume of the region in the respective group.

The brain–behavior relationships were tested in trained mice scanned four times (*n* = 17) by correlating the change from S1 to S2 in the regional volume, expressed as a fraction of total brain volume (V_S2_/TBV_S2_ − V_S1_/TBV_S1_), with reaching performance defined as the mean success rate across the two full training sessions flanking scan S2 (training days 2 and 4). Pearson correlation with Fisher-*z* confidence intervals was the primary analysis and Spearman rank correlation (tied scores given average ranks; *p* values and confidence intervals based on the Bonnett-Wright standard error) a sensitivity analysis, each Holm-corrected across the two regions separately. To test whether the associations reflected global volume change, the raw S1 to S2 change in total brain volume (TBV_S2_ − TBV_S1_, in mm^3^) of the same mice was correlated with the same performance measures (Pearson correlation with Fisher-z confidence interval, not corrected for multiple comparisons).

#### Humans

MRI data analyses were performed using Matlab 2023a, the SPM12 toolbox (version 7771), the Computational Anatomy Toolbox (CAT12; version CAT12.8.2 r2159) and the hMRI toolbox (version 0.6.1). T_1_-weighted anatomical images were preprocessed using CAT12 with the longitudinal processing pipeline using the neuroplasticity model with default parameters. This pipeline is optimized for detecting subtle within-participant anatomical changes over time. For each participant, images from all timepoints were realigned and corrected for intra-subject bias-field inhomogeneities. A high-quality within-subject average T_1_-weighted image was then created across all available timepoints. The average image was segmented to generate subject-specific tissue probability maps. Using these priors, each individual timepoint was segmented. The resulting tissue segments were spatially normalized to MNI space using an average of the timepoint-specific registration parameters. Normalized gray matter tissue segments were modulated and subsequently smoothed with an isotropic Gaussian kernel of 6 mm full-width at half maximum (FWHM).

Quantitative multi-parameter maps (MT_sat_, PD, R_1_, and R_2_*) were generated for each session using the default parameters of the hMRI toolbox. All available sessions from each participant were included in the preprocessing. For spatial processing, we adapted the standard hMRI preprocessing pipeline for longitudinal analysis, following the protocol described by Wenger et al. (2022). Multichannel tissue segmentation was performed for each session using the synthetic MT-weighted (MTw), proton density-weighted (PDw), and T_1_-weighted (T_1_w) images, which were computationally generated from the MPM protocol to provide complementary tissue contrast. Tissue probability maps for gray matter, white matter, and cerebrospinal fluid were derived from these synthetic images using tissue probability maps based on multiparametric data (eTPMs) to improve segmentation accuracy for quantitative MRI (Draganski et al., 2011).

Longitudinal image registration was conducted using SPM12’s Geodesic Shooting (SHOOT) algorithm. First, within-subject templates were created by registering all sessions for each participant, providing an unbiased average structural reference. Subsequently, a group (between-subject) template was generated by registering all within-subject templates using the same SHOOT approach. The deformation fields from both template creation steps (i.e., from each session’s native space to its within-subject template, and from the within-subject template to the group template) were combined and used to normalize all quantitative parameter maps from each session to group template space and, finally, to MNI space. All normalized parameter maps were smoothed using tissue-weighted smoothing (Draganski et al., 2011) with a 6 mm FWHM Gaussian kernel prior to statistical analysis.

For the CAT12-derived morphometric data, we used general linear models with a flexible factorial design including three factors: Subject, Time (with up to seven available time points per participant), and Group (training versus control). This design was used to test Time × Group interactions, assessing whether the training and control group differed in predefined longitudinal patterns of change (*n* = 36 training, *n* = 35 control). Across all seven time points, we tested a curvilinear interaction contrast reflecting an increase toward the middle of the training period, peaking around S4, followed by a subsequent decrease toward S7. This contrast used the temporal weights [−1.00, −0.05, 0.60, 1.00, 0.65, −0.20, −1.00] for the training group and the opposite weights for the control group. In addition, linear and quadratic trajectories across all time points were tested. Further exploratory analyses, including analyses informed by individual learning curves and ROI analyses based on task-related activation are described in the Supplementary Methods. Motivated by findings of rapid structural plasticity, we also performed an exploratory pairwise interaction analysis comparing change from S1 to S2 between groups, testing both directions. Results were thresholded at a voxel-wise threshold of *p* < .001 uncorrected, with a minimum cluster extent of k = 50 voxels.

For the hMRI-derived qMRI maps, we used a flexible factorial model with factors Subject, Time, and Group. Motivated by rapid plasticity, the regional volume findings early in learning I mice, and the absence of detectable VBM-based gray matter volume changes in humans, we performed a Time × Group interaction contrast between S1 and S2 to assess whether qMRI parameters showed early training-related differences between groups (*n* = 33 training, *n* = 30 control). We used an explicit mask covering the whole brain to restrict the search volume. For all qMRI parameter maps, a cluster-size threshold for statistical significance was determined using AFNI’s 3dClustSim. Based on the simulation table, a minimum cluster extent of 232 voxels was used, corresponding to *p* < .001 with a corrected alpha of .05. Therefore, results in the maps were considered significant at a voxel-wise threshold of *p* < .001 uncorrected and a cluster-extent threshold of k ≥ 232 voxels. For each significant cluster, mean qMRI values were extracted for each participant, and the change in qMRI measure between S1 and S2 was correlated with behavioral performance on the chopstick grasping task at S2 using Pearson correlations.

## Data Availability

Anonymized human and mouse behavioral data, human MRI-derived data extracted from the significant clusters identified in the whole-brain analysis, and mouse MRI data underlying the ROI analyses will be deposited in a public repository upon publication. Raw human MRI data (i.e., individual brain images) cannot be made publicly available because of restrictions related to participant consent, ethical approval, and applicable data protection requirements. Analysis code is available from the corresponding author upon request and will be made publicly available on GitHub upon publication.

## CRediT author statement

**Maike Hille**: Data curation, Formal analysis, Investigation, Methodology, Visualization, Writing – original draft, Writing – review & editing. **Lena C. Justus:** Formal analysis, Writing – original draft, Writing – review & editing. **Sara Zocher:** Data curation, Investigation, Methodology, Writing – review & editing. **Tibor Stark:** Data curation, Investigation, Methodology, Writing – review & editing. **Moritz Bonhoeffer:** Formal analysis, Writing – review & editing. **Jason P. Lerch:** Methodology, Formal analysis, Writing – review & editing. **Michael Czisch:** Data curation, Investigation, Methodology, Writing – review & editing. **Simone Kühn:** Conceptualization, Methodology, Resources, Supervision, Writing – review & editing. **Ulman Lindenberger:** Conceptualization, Methodology, Resources, Supervision, Writing – review & editing. **Tobias Bonhoeffer:** Conceptualization, Methodology, Resources, Supervision, Writing – review & editing.

## Declaration of generative AI and AI-assisted technologies in the manuscript preparation process

During the preparation of this work, the authors used ChatGPT (OpenAI; GPT-5.6) and Claude (Anthropic; Opus 5.5) to support implementation of statistical analysis approaches and to refine and polish the structure, formatting, and language of the manuscript. After using these tools, the authors reviewed and edited the content as needed and take full responsibility for the content of the published article.

## Acknowledgments

We acknowledge the financial support of the Max Planck Society. This work was conducted at the Max Planck Dahlem Campus of Cognition (MPDCC) of the Max Planck Institute for Human Development, Berlin, Germany, at the Max Planck Institute for Biological Intelligence (former MPI of Neurobiology), Martinsried, Germany, and at the Max Planck Institute of Psychiatry, Munich, Germany. During the work on her dissertation, M.H. was a pre-doctoral fellow of the International Max Planck Research School on Learning, Institutions, and Future Evolution (LIFE). The authors thank Carsten Wotjak, Elisabeth Binder, and Nadine Gogolla for helpful discussions, valuable feedback, and intellectual contributions during the development of the project. We thank Sebastian Schröder, Manuel Teichert, Volker Staigler, and the workshop of the MPI of Psychiatry for designing and constructing the custom experimental equipment used in this study. We thank Stefan Hetzer for assistance with the acquisition and preprocessing of the quantitative MRI multiparameter mapping protocol. We also thank Julia Delius for editorial assistance.

