## Supplementary Material for "Investigating Motor Skill Acquisition in Mice and Humans: Towards A Cross-Species Paradigm"

### *Methods*

### <u>Analysis of CAT12-derived morphometric data</u>

In addition to the standard longitudinal flexible factorial analysis, we performed an exploratory analysis in which the model was informed by each participant's individual behavioral learning curve. For each participant, daily home-training performance was modeled as a function of training day in R using the segmented package (https://cran.r-project.org/web/packages/segmented/index.html). Specifically, we fitted individual piecewise linear regression models to the behavioral learning curves to estimate a breakpoint, defined as the training day on which the slope of the learning curve changed. The estimated behavioral breakpoint was then assigned to the MRI time point closest in time to that training day, and participants were classified accordingly. In SPM12, for each participant, we constructed a participant-specific temporal contrast vector across the seven MRI sessions. This vector modeled an increase in gray matter volume up to the MRI session closest to the behavioral breakpoint, followed by a decrease after the breakpoint. The vectors were scaled to a maximum absolute value of 1 to make contrasts comparable across participants. The resulting contrast images were then entered into a second-level one-sample t-test, testing for common significant voxels for the training group. This analysis included 33 participants from the training group who completed the study.

We also performed a region-of-interest (ROI) analysis using activation clusters from the two fMRI tasks. For the finger tapping task and the chopstick grasping task, first-level contrast images were computed for right finger tapping versus baseline and chopstick grasping versus baseline, respectively. These contrast images were separately entered into second-level one-sample t-tests across participants to identify brain regions significantly activated by each condition. To restrict the analysis to relevant motor regions, significant activation clusters were masked with the precentral gyrus, postcentral gyrus and supplementary motor area ROIs from the AAL3 atlas. This resulted in task-specific ROIs from which we extracted the mean gray matter volume values from the CAT12-preprocessed structural images for each participant and MRI timepoint. The extracted values were analyzed in R using linear mixed-effects models to test whether structural gray matter estimates within the functionally defined ROIs changed over time differently in each group. Specifically, linear mixed-effects models were fitted in R using gray matter volume estimates extracted from each ROI as the dependent variable. Fixed effects included time, group, and the Time × Group interaction. Participant was included as a random intercept to account for repeated measurements. The main effect of interest was the Time × Group interaction, indicating whether gray matter volume changed differently between the groups within the functionally defined ROI.

To obtain a more anatomically individualized estimate of gray matter volume change in the primary motor cortex hand area, we performed an additional ROI analysis based on participant-specific functional activation peaks. For each participant, the peak activation voxel for the contrast right finger tapping versus baseline was identified in the primary motor cortex hand region. Around this individual peak coordinate, a sphere with a radius of 6 mm was created. Mean gray matter volume estimates from the CAT12-preprocessed structural images were then extracted from these individually defined spherical ROIs for each participant and MRI timepoint. The extracted gray matter volume estimates were analyzed in R using linear mixed-effects models with time, group, and their interaction as fixed effects and participant as a random effect. The effect of interest was the Time × Group interaction, testing whether longitudinal gray matter volume changes in the individually defined primary motor cortex hand region differed between groups.

**Analyses of task-based functional MRI data**

Functional MRI data were preprocessed in SPM12 using default parameters, including slice time correction, realignment, co-registration to the structural image, normalization to MNI space, and spatial smoothing using an 8-mm full-width at half-maximum (FWHM) Gaussian kernel. First-level statistical analyses were performed using the general linear model in SPM12. Task conditions were modeled and convolved with the canonical hemodynamic response function, with motion parameters included as nuisance regressors. Contrasts were computed for right finger tapping versus baseline, left finger tapping versus baseline in the finger tapping task, and chopstick grasping versus baseline for the chopstick grasping task. These contrast images were then entered into second-level one-sample t-tests to identify task-related activation across participants.

*Results*

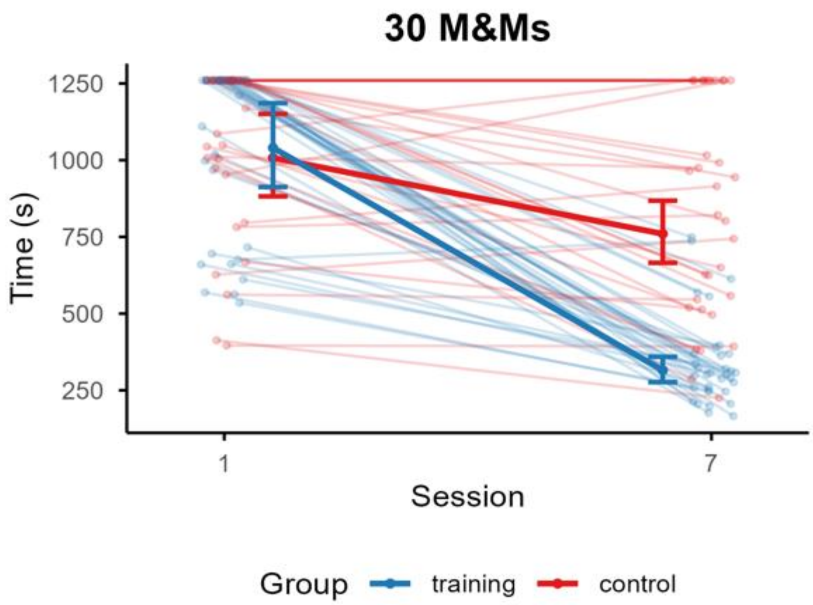


**Figure S1.** Performance on the trained chopstick grasping task before (S1) and after (S7) training in the chopstick training ($n$ = 33) and reading control ($n$ = 30) groups. Individual points and lines show participant-level performance across sessions. Bold points and lines show model-estimated marginal means, with error bars indicating 95% confidence intervals.

**Table S1**

*Behavioral performance on chopstick grasping tasks*

| Statistic | 30 M&Ms | 30 Maltesers | 10 Marbles | 10 M&Ms (left hand) | Purdue Pegboard with chopsticks |
|---|---|---|---|---|---|
| **Session main effect** | $F$(1, 63) = 176.14, $\mathbf{p < .001}$ | $F$(1, 65.4) = 62.59, $\mathbf{p < .001}$ | $\chi^2(1) = 20.94$, $\mathbf{p < .001}$ | $\chi^2(1) = 21.46$, $\mathbf{p < .001}$ | $\chi^2(1) = 0.19$, $p = .666$ |
| **Group main effect** | $F$(1, 63) = 31.10, $\mathbf{p < .001}$ | $F$(1, 67.2) = 2.19, $p = .143$ | $\chi^2(1) = 0.68$, $p = .409$ | $\chi^2(1) = 0.40$, $p = .528$ | $\chi^2(1) = 0.38$, $p = .536$ |
| **Group × Session interaction** | $F$(1, 63) = 67.27, $\mathbf{p < .001}$ | $F$ (1, 65.4) = 33.29, $\mathbf{p < .001}$ | $\chi^2(1) = 6.07$, $\mathbf{p = .014}$ | $\chi^2(1) = 11.15$, $\mathbf{p < .001}$ | $\chi^2(1) = 5.11$, $\mathbf{p = .024}$ |
| **Chopstick: S1 → S7** | 1040 s → 315 s −69.7% ($t$(63) = −15.30, $\mathbf{p < .001}$) | 260 s → 125 s −51.8% ($t$(66.4) = −9.74, $\mathbf{p < .001}$) | 20.0% → 72.2% success ($z$ = 4.89, $\mathbf{p < .001}$) | 41.2% → 90.0% success ($z$ = 5.52, $\mathbf{p < .001}$) | 3.77 → 4.92 pegs +30.6% ($z$ = 1.93, $p = .054$) |

| Statistic | 30 M&Ms | 30 Maltesers | 10 Marbles | 10 M&Ms (left hand) | Purdue Pegboard with chopsticks |
|---|---|---|---|---|---|
| **Reading: S1 → S7** | 1008 s → 760 s; −24.6% ($t(63) = -3.56$, $\mathbf{p < .001}$) | 212 s → 189 s; −10.8% ($t(66.5) = -1.50$, $p = .138$) | 26.9% → 47.0% success ($z = 2.21$, $\mathbf{p = .027}$) | 53.4% → 67.9% success ($z = 1.66$, $p = .097$) | 4.31 → 3.66 pegs −15.2% ($z = -1.25$, $p = .212$) |
| **Group difference at S1** | n.s. ($t(115) = 0.34$, $p = .735$) | chopstick group 22.6% slower ($t(123) = 2.36$, $\mathbf{p = .020}$) | n.s. chopstick −7.0 pp ($z = -0.88$, $p = .378$) | n.s.; chopstick −12.2 pp ($z = -1.16$, $p = .248$) | n.s.; chopstick 12.7% lower ($z = -0.64$, $p = .523$) |
| **Group difference at S7** | chopstick group 58.5% faster; ($t(115) = -9.34$, $\mathbf{p < .001}$) | chopstick group 33.7% faster ($t(126) = -4.57$, $\mathbf{p < .001}$) | chopstick +25.2 pp ($z = 2.38$, $\mathbf{p = .017}$) | chopstick +22.1 pp ($z = 2.78$, $\mathbf{p = .006}$) | n.s.; chopstick 34.5% higher ($z = -1.49$, $p = .137$) |

*Note.* Statistical results for the chopstick training and reading control groups across the first (S1) and final (S7) sessions on all chopstick grasping tasks. pp = percentage points; s = seconds. Bold *p* values indicate statistically significant results ($p < .05$).

**Table S2**

*Behavioral performance on general fine motor dexterity tasks*

| Statistic | Purdue Pegboard | | | |
|---|---|---|---|---|
| | **left** | **right** | **both hands** | **assembly** |
| **Session main effect** | $F(1, 64.6) = 10.07$, $\mathbf{p = .002}$ | $F(1, 63.0) = 16.68$, $\mathbf{p < .001}$ | $F(1, 67.2) = 12.05$, $\mathbf{p < .001}$ | $F(1, 64.8) = 8.13$, $\mathbf{p = .006}$ |
| **Group main effect** | $F(1, 68.7) = 0.02$, $p =.875$ | $F(1, 66.2) = 0.002$, $p = .965$ | $F(1, 70.2) = 0.13$, $p = .719$ | $F(1, 67.5) = 2.84$, $p = .096$ |
| **Group × Session interaction** | $F(1, 64.6) = 0.48$, $p = .492$ | $F(1, 63.0) = 0.02$, $p = .899$ | $F(1, 67.2) = 0.001$, $p = .970$ | $F(1, 64.8) = 0.01$, $p = .906$ |
| **Group difference at S1** | n.s. ($t(94.6) = 0.45$, $p = .658$) | n.s. ($t(106) = 0.10$, $p = .918$) | n.s. ($t(109) = -0.33$, $p = .740$) | n.s. ($t(113) = -1.38$, $p = .171$) |

| Statistic | Grooved Pegboard | | O'Connor |
|---|---|---|---|
| | **left** | **right** | |
| **Session main effect** | $F(1, 63.0) = 15.19$, $\boldsymbol{p < .001}$ | $F(1, 63.5) = 11.66$, $\boldsymbol{p = .001}$ | $F(1, 65.8) = 0.98$, $p = .326$ |
| **Group main effect** | $F(1, 67.2) = 0.02$, $p = .896$ | $F(1, 67.3) = 0.11$, $p = .746$ | $F(1, 68.8) = 2.69$, $p = .105$ |
| **Group × Session interaction** | $F(1, 63.0) = 0.78$, $p = .381$ | $F(1, 63.5) = 0.01$, $p = .909$ | $F(1, 65.8) = 0.05$, $p = .831$ |
| **Group difference at S1** | n.s. ($t(92.7) = 0.25$, $p = .804$) | n.s. ($t(99) = -0.24$, $p = .811$) | n.s. ($t(109) = -1.31$, $p = .193$) |

*Note*. Statistical results for the chopstick training and reading control groups across the first (S1) and final (S7) sessions on all general fine motor dexterity tasks. Bold $p$ values indicate statistically significant results ($p < .05$).

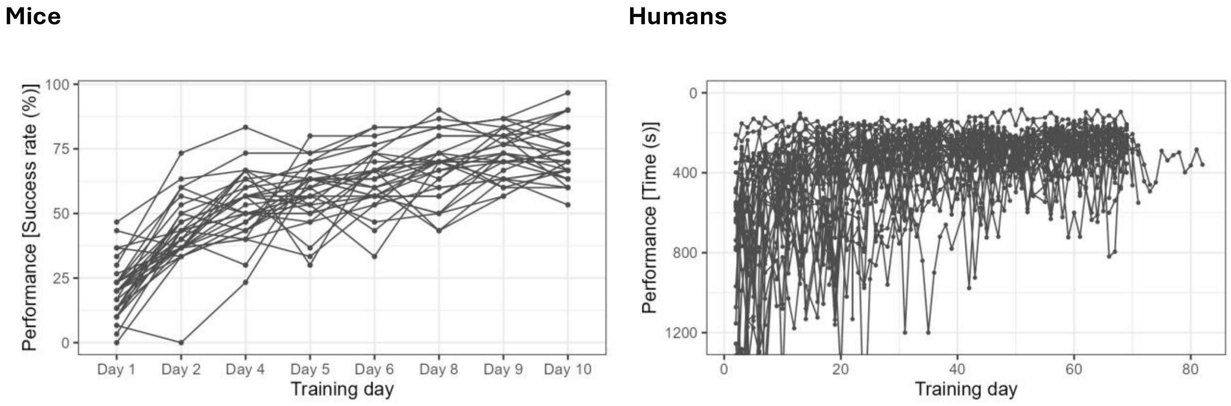


**Figure S2.** Individual learning curves for trained mice (right, $n = 32$) and humans (left, $n = 36$) across training sessions. Each line represents one individual and points indicate performance on a given day. Mouse performance is expressed as success rate in percent, with higher values indicating better performance. Human performance is expressed as completion time in seconds, with lower values indicating better performance; the y-axis is reversed so that improvement is displayed upward.

**Table S3**

*Regional volume changes from S1 to S2 in mice*

| Region | Group | Change (%) | Estimate (mm$^3$) | 95% CI (mm$^3$) | $t$(df) | $p$ | $p_{Holm}$ |
|---|---|---|---|---|---|---|---|
| **Primary motor** | training | −1.3 | −0.046 | [−0.095, 0.003] | −1.85 (90.4) | .067 | .20 |
| **Primary motor** | control | 0.7 | 0.025 | [−0.026, 0.007] | 0.98 (95.8) | .33 | .66 |
| **Forelimb somatosensory** | training | −2.5 | −0.048 | [−0.072, -0.024] | −3.96 (96.5) | **< .001** | **<.001** |
| **Forelimb somatosensory** | control | −0.1 | −0.002 | [−0.028, 0.023] | −0.17 (101.1) | .86 | .86 |

*Note.* Regional volume changes from the S1 (day −3) to S2 (day 3) within each group, estimated from the linear mixed model (all four scans) with total brain volume as a covariate. Percentages are relative to each group's own mean baseline volume of the region. $t$ tests with Satterthwaite-approximated degrees of freedom; $p_{\text{Holm}}$, Holm-corrected across the four tests. Control $n$ = 16 mice, trained $n$ = 17 mice.

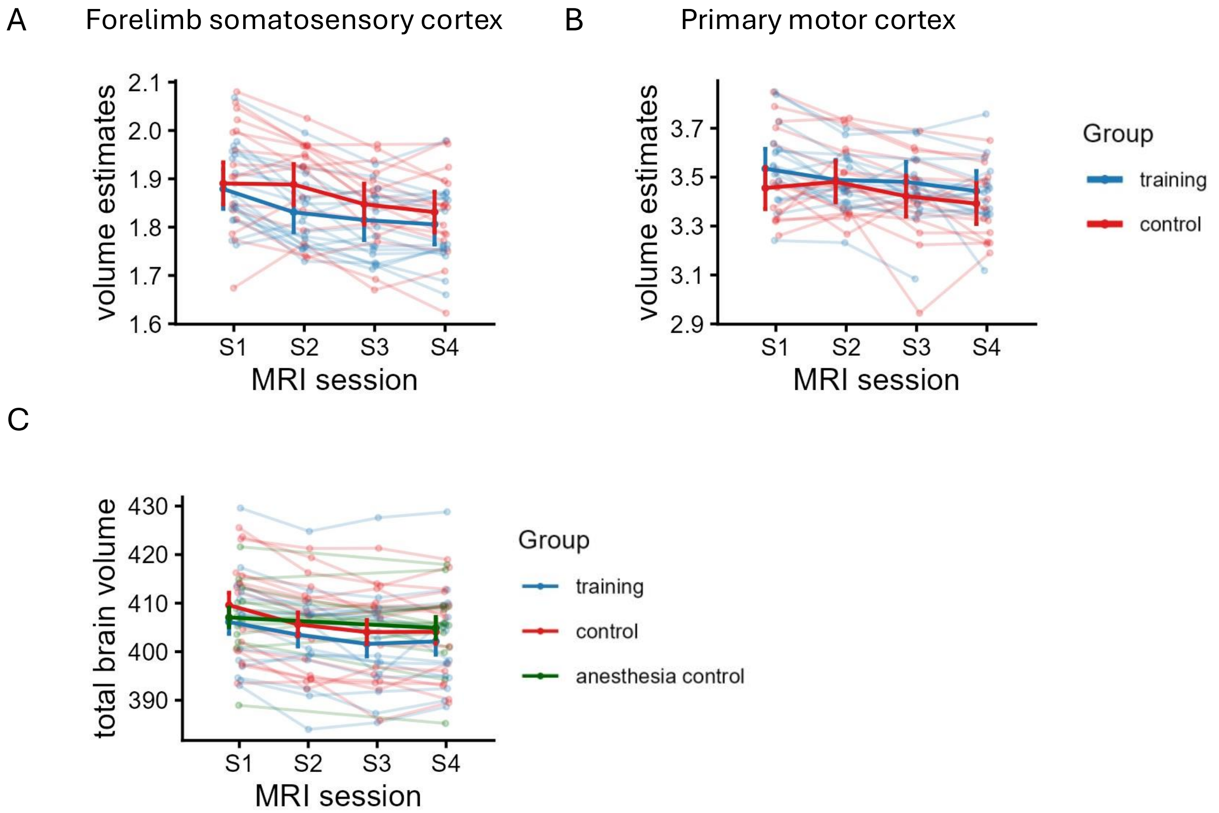


**Figure S3.** ***A,*** Regional brain volume for the forelimb somatosensory cortex across all four MRI sessions. ***B,*** Regional brain volume for the primary motor cortex across all four MRI sessions. ***C,***

Total brain volume over the complete imaging period for the three groups of mice. Control mice $n = 16$, trained mice $n = 17$ (minus one scan in session 4), anesthesia control $n = 13$.

**Table S4**

*Baseline comparison of quantitative MRI parameters*

| **Map** | **Cluster** | **Mean training group** | **Mean control group** | ***t*** | **df** | ***p*** | **Cohen's *d*** |
|---|---|---|---|---|---|---|---|
| **$MT_{sat}$** | x = −50, y = 3, z = 51 | 0.894 | 0.894 | −0.017 | 63.9 | .987 | −0.004 |
| **$MT_{sat}$** | x = −31, y = 59, z = 16 | 0.858 | 0.854 | 0.151 | 60.6 | .880 | 0.037 |
| **PD** | x = −66, y = −15, z = −31 | 78.9 | 79.9 | −2.38 | 56.5 | **.021** | −0.585 |
| **PD** | x = −21, y = 1, z = 66 | 79.4 | 78.7 | 1.49 | 64.0 | .142 | 0.366 |
| **$R_1$** | x = −35, y = 12, z = 34 | 0.630 | 0.643 | −1.72 | 63.6 | .090 | −0.423 |
| **$R_2^*$** | x = −12, y = −69, z = −56 | 19.3 | 18.0 | 1.84 | 63.2 | .070 | 0.454 |
| **$R_2^*$** | x = −26, y = −74, z = −52 | 17.8 | 17.0 | 3.38 | 64.0 | **.001** | 0.832 |

*Note.* Baseline (S1) comparison of quantitative MRI measures between training and control groups.

**Table S5**

*Pearson correlations between qMRI parameter changes and behavioral performance*

| **Map** | **Cluster** | **Pearson r** | ***p*** |
|---|---|---|---|
| **$MT_{sat}$** | x = −50, y = 3, z = 51 | −0.135 | .463 |
| **$MT_{sat}$** | x = −31, y = 59, z = 16 | −0.208 | .254 |
| **PD** | x = −21, y = 1, z = 66 | −0.123 | .504 |
| **$R_1$** | x = −35, y = 12, z = 34 | 0.013 | .944 |
| **$R_2^*$** | x = −12, y = −69, z = −56 | −0.191 | .296 |

*Note.* Pearson correlations between qMRI parameter changes from S1 to S2 and behavioral performance at S2.

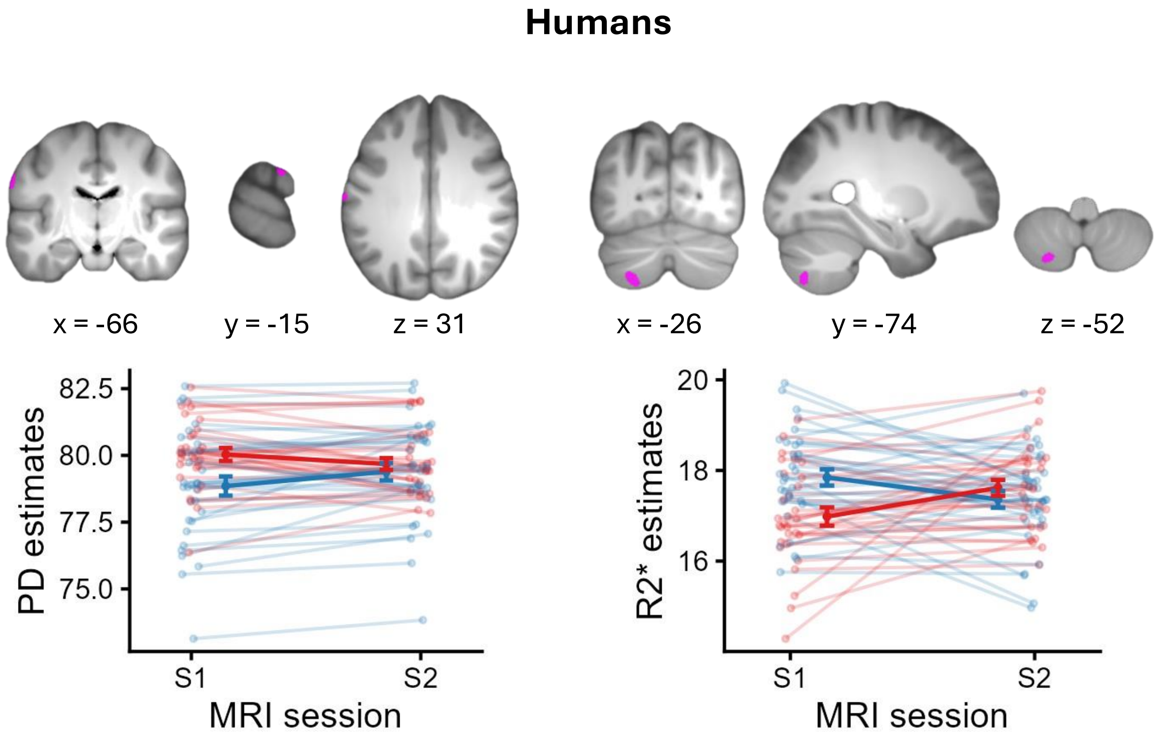


**Figure S4.** Changes in qMRI parameter maps from session 1 to session 2 in humans ($p < .001$ uncorrected, cluster-size threshold k = 232 voxels).
